# Oil body formation in *Marchantia polymorpha* is controlled by MpC1HDZ and serves as a defense against arthropod herbivores

**DOI:** 10.1101/2020.03.02.971010

**Authors:** Facundo Romani, Elizabeta Banic, Stevie N. Florent, Takehiko Kanazawa, Jason Q.D. Goodger, Remco Mentink, Tom Dierschke, Sabine Zachgo, Takashi Ueda, John L. Bowman, Miltos Tsiantis, Javier E. Moreno

## Abstract

The origin of a terrestrial flora in the Ordovician required adaptation to novel biotic and abiotic stressors. Oil bodies, a synapomorphy of liverworts, accumulate secondary metabolites, but their function and development are poorly understood. Oil bodies of *Marchantia polymorpha* develop within specialized cells as one single large organelle. Here, we show that a CLASS I HOMEODOMAIN LEUCINE-ZIPPER (C1HDZ) transcription factor controls the differentiation of oil body cells in two different ecotypes of the liverwort *M. polymorpha*, a model genetic system for early divergent land plants. In flowering plants, these transcription factors primarily modulate responses to abiotic stresss including drought. However, loss-of-function alleles of the single ortholog gene, Mp*C1HDZ*, in *M. polymorpha* did not exhibit phenotypes associated with abiotic stress. Rather Mp*c1hdz* mutant plants were more susceptible to herbivory and total plant extracts of the mutant exhibited reduced antibacterial activity. Transcriptomic analysis of the mutant revealed a reduction in expression of genes related to secondary metabolism that was accompanied by a specific depletion of oil body terpenoid compounds. Through time lapse imaging we observed that MpC1HDZ expression maxima precede oil body formation indicating that MpC1HDZ mediates differentiation of oil body cells. Our results indicate that *M. polymorpha* oil bodies, and MpC1HDZ, are critical for defense against herbivory but not for abiotic stress-tolerance. Thus, C1HDZ genes were co-opted to regulate separate responses to biotic and abiotic stressors in two distinct land plant lineages.

## INTRODUCTION

The emergence of land flora in the Ordovician and the subsequent radiation of plants into terrestrial environments were major evolutionary events in the history of Earth. This radical breakthrough for life on the planet entailed the evolution of both new developmental programs and biochemical pathways [1]. Early land plants were faced with novel biotic and abiotic stressors, including a desiccating aerial environment, increased solar radiation, and greater interspecific competition, including herbivory. This transition was accompanied by a dramatic increase in the diversity of secondary metabolites and the complexity of transcriptional regulation, to protect plants from herbivores and pathogens, and also abiotic stressors such as desiccation and light intensity [2-4].

Liverworts represent one of the earliest diverging land plant lineages, with a predicted Ordovician-Silurian origin of all three major extant liverwort clades [5, 6]. One synapomorphy of this lineage is the presence of oil bodies, specialized organelles containing an array of lipophilic secondary compounds [7]. In some liverworts, multiple oil bodies are found in nearly all differentiated cells of both gametophyte and sporophyte generations. Whilst in other liverworts, such as *Marchantia polymorpha* (*M. polymorpha*), oil bodies are large and differentiate in only a subset of isolated cells referred to as idioblasts. They originate in young meristematic cells in a relatively constant proportion [8]. It has long been known that the liverwort fragrance is associated to the oil body presence [9], and that species-specific differences in taste correlate with oil body chemical composition [10]. Early biochemical analyses identified numerous terpenoids as major chemical constituents of liverwort oil bodies [11, 12], and later studies expanded this list to include diterpenes, bis(bibenzyls), and additional compounds [13-16]. Pfeffer, who gave oil bodies their name (*Ölkörper*), noted they were membrane bound [17], and later studies confirmed a single membrane [18, 19] in which isoprenoid biosynthesis enzymes are localized [16].

Oil body function remains contested, with suggested defensive roles against both abiotic and biotic stressors. Roles protecting against a wide variety of abiotic stressors have been proposed, including cold and osmotic stress [7], desiccation [20-22], high light intensity [23], UV radiation [24, 25], and metabolic stress [18]. In addition, a role against herbivory was postulated by Stahl, who presented snails with fresh or alcohol-leached liverworts and noted that snails preferred the latter and did not consume the former to any extent [26]. An anti-herbivory role is also supported by fossilized evidence of herbivore oil body avoidance in Devonian liverworts [27]. Furthermore, the chemical constituents of oil bodies lead to growth inhibition of bacterial and fungal pathogens [28-32]. Despite a large body of literature describing the presence and diversity of oil body secondary metabolites [14, 15, 33, 34], less is known of the regulatory circuits leading to oil body cell differentiation.

Synthesis of specialized metabolites in vascular plants is tightly regulated by transcription factors (TFs), with some orthologs having conserved and others novel roles in liverworts [32, 35-38]. Class I HOMEODOMAIN LEUCINE-ZIPPER (C1HDZ) TFs genes are conserved across streptophytes with a single homolog predicted to have been present in the common-ancestor of land plants [39]. In angiosperms, C1HDZ genes are well-known regulators of tolerance to abiotic stress [40, 41], but also control other molecular functions including leaf, root and silique development and vascular patterning [42-44]. However, little is known about the physiological role of C1HDZ in bryophytes [45]. *M. polymorpha* provides a model to investigate their functional evolution since the genome encodes only one single C1HDZ ortholog (Mp*C1HDZ*) [2]. Here, we investigate Mp*C1HDZ* functionality and report that it exhibits a divergent function relative to its angiosperms orthologs.

## RESULTS

### Mp*C1HDZ* controls the development of oil body cells

Mp*C1HDZ*, the solitary C1HDZ *M. polymorpha* ortholog, possesses all conserved protein motifs [39] and predicted structural features previously defined for C1HDZ proteins (Figure 1A, Figure S1A). MpC1HDZ exhibits nuclear localization in transient expression assays in tobacco plants (Figure S1B). Using two different genetic backgrounds (WT = Australian, BoGa = Botanical Garden Osnabrück, German accession), we targeted gRNAs to the HDZ domain generating mutant alleles with premature STOP codons, deletion of the whole HDZ domain or deletion of the entire *locus* (Figure 1A). Four lines were clonally purified through gemmae generations: Mp*c1hdz-1*^*ge*^ (male) and Mp*c1hdz-2*^*ge*^ (female) harbor frameshift mutations that generate early stop codons and were generated in the Australian accession. Mp*c1hdz-3*^*ge*^ and Mp*c1hdz-4*^*ge*^ were generated in the BoGa accession deleting the whole CDS or the HDZ domain, respectively (Figure 1A; nomenclature as in [46]).

**Figure 1.**
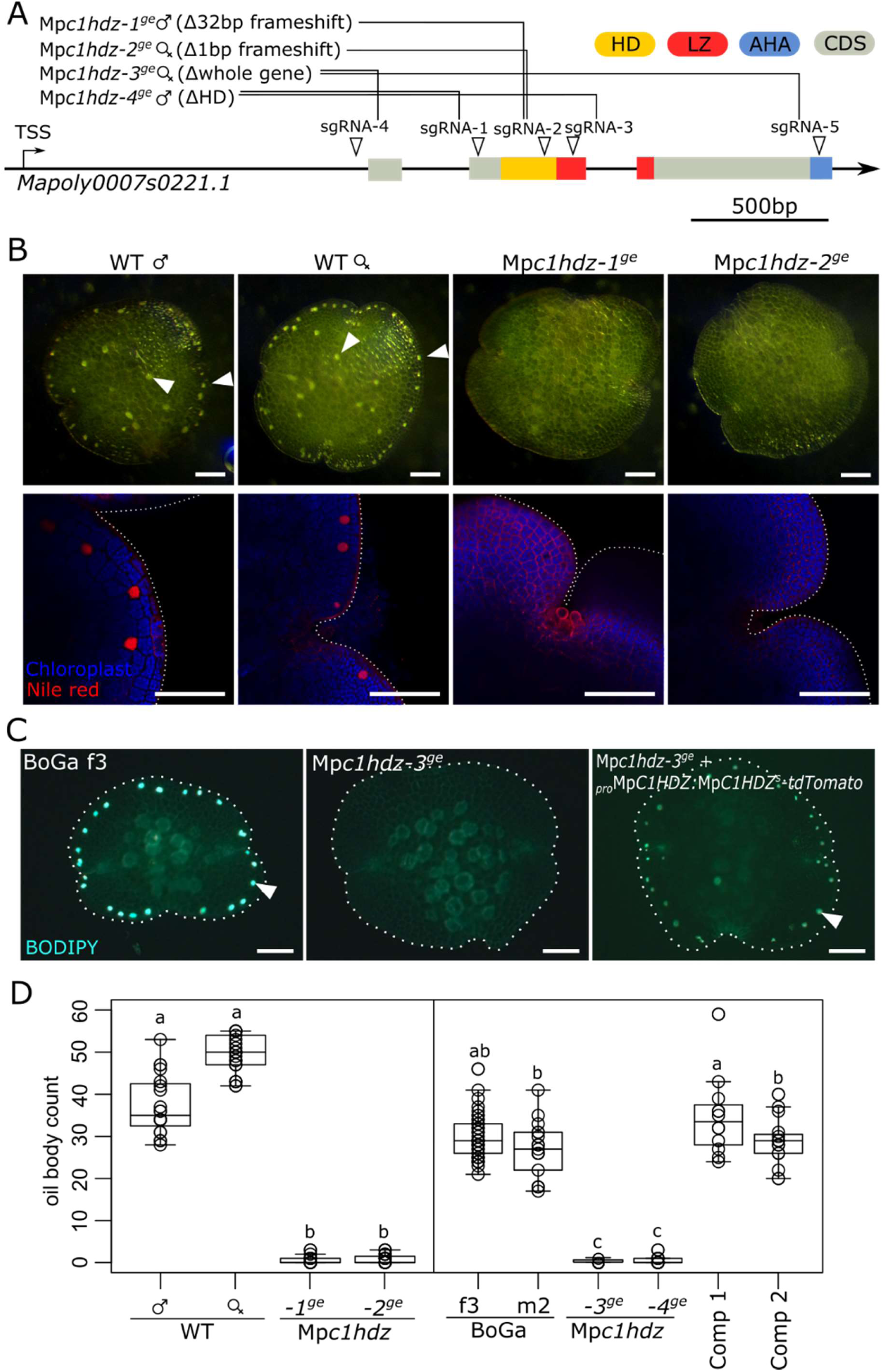
Oil body number is strongly reduced in Mp*c1hdz* plants. (A) Schematic representation of the Mp*C1HDZ* gene domains and location of designed CRISPR gRNAs targeting the genomic locus (arrowheads). Colored boxes represent the three conserved protein domains. The combination of gRNAs used to generate frameshifts deletions in each case is shown. (B) Phenotype of freshly collected gemmae from wild-type and Mp*c1hdz* mutant plants under light microscopy (upper panel); confocal microscopy of Red Nile stained gemmae (bottom, panel). Red Nile fluorescence is shown in red and chloroplast auto-fluorescence in blue. (C) Oil body formation is restored in gemmae expressing Mp*C1HDZ* translational reporter _*pro*_Mp*C1HDZ*^*3kb*^:Mp*C1HDZ-tdTomato* and visualized by BODIPY staining. (D) Number of oil bodies per gemma in wild-type, mutant and complemented plants in both genetic backgrounds were quantified and represented as boxplot (n>12). Comp 1: Mp*c1hdz*-3^ge^ + _*pro*_Mp*C1HDZ*^*3kb*^:Mp*C1HDZ-Citrine*; Comp 2: Mp*c1hdz*-3^ge^ + _*pro*_Mp*C1HDZ*^*3kb*^:Mp*C1HDZ-tdTomato*. Statistical difference was tested using ANOVA followed by Tukey HSD *p* value < 0.01; letters indicate statistically significant groups.

In all cases, mutant plants did not show gross phenotypic defects and completed their life cycle in a similar manner and timing as wild type (Figure S2). To understand the physiological role of Mp*C1HDZ*, we performed a detailed phenotypic characterization of the mutant alleles. The most conspicuous phenotype of Mp*c1hdz* mutants is a striking reduction in the number of oil bodies, both in gemmae and thalli, in two different genetic backgrounds (Figure 1B,C). *M. polymorpha* oil body cells are observed as brown dots or highly-reflective cells, with 20-50 oil bodies per gemmae, mostly located marginally (Figure 1B,C). Whereas mutant gemmae lacked oil bodies, in the thallus and in the parenchymatic region we found a small number of oil bodies in the tissue (Figure S3). It is known that specific environmental conditions, such as starvation and field non-axenic cultivation, affect oil body cell number [15]. Wild type plants grown in Petri-dishes developed fewer oil body cells compared to non-axenic grown plants and starvation-treated plants (Figure S3). In contrast, the number of oil body cells of the Mp*c1hdz* mutant remained low in both conditions (Figure S3).

To examine oil body distribution throughout the life cycle, we crossed Mp*c1hdz-2*^*ge*^ with Mp*c1hdz-1*^*ge*^. The resultant sporophyte produced spores of the two genotypes and sporelings were phenotypically wild type. After 18-days, oil bodies were visible in wild type sporelings, but were totally absent in Mp*c1hdz* sporelings (Figure S3C). Thus, we conclude that MpC1HDZ is a positive regulator of oil body formation across all organs tested and developmental stages.

To confirm the role of Mp*C1HDZ* in the oil body phenotype, we complemented the mutant using the endogenous promoter of Mp*C1HDZ* driving Mp*C1HDZ*-*tdTomato* (_*pro*_Mp*C1HDZ*^*3kb*^:Mp*C1HDZ*-*tdTomato*) in the BoGa ecotype, which rescued the oil body phenotype (Figure 1C-D). In contrast, when Mp*C1HDZ* expression was driven by either of two constitutive promoters (_*pro*_*EF1:*Mp*C1HDZ*, _*pro*_*35S:*Mp*C1HDZ*) no viable plants were obtained (Figure S4), suggesting that constitutive overexpression is lethal (data not shown).

### The Mp*C1HDZ* promoter is active in thallus and gemmalings with high expression in oil body cells

To better understand the relationship between Mp*C1HDZ* and oil body cells, we studied the gene expression pattern with transcriptional reporters (_*pro*_Mp*C1HDZ:GUS*) using the putative regulatory region of each genetic background. In 3-4 day-old gemmalings, strong GUS activity was observed in oil body cells (Figure 2A). At later stages, GUS activity was detected in the center of the thallus and in meristematic regions (Figure 2B,C), at the notch extending into the midrib, including the gemma cups (Figure 2C-E), and broadly in reproductive organs (Figure 2F). A similar transcriptional reporter activity was also observed in the BoGa ecotype (Figure S5).

**Figure 2.**
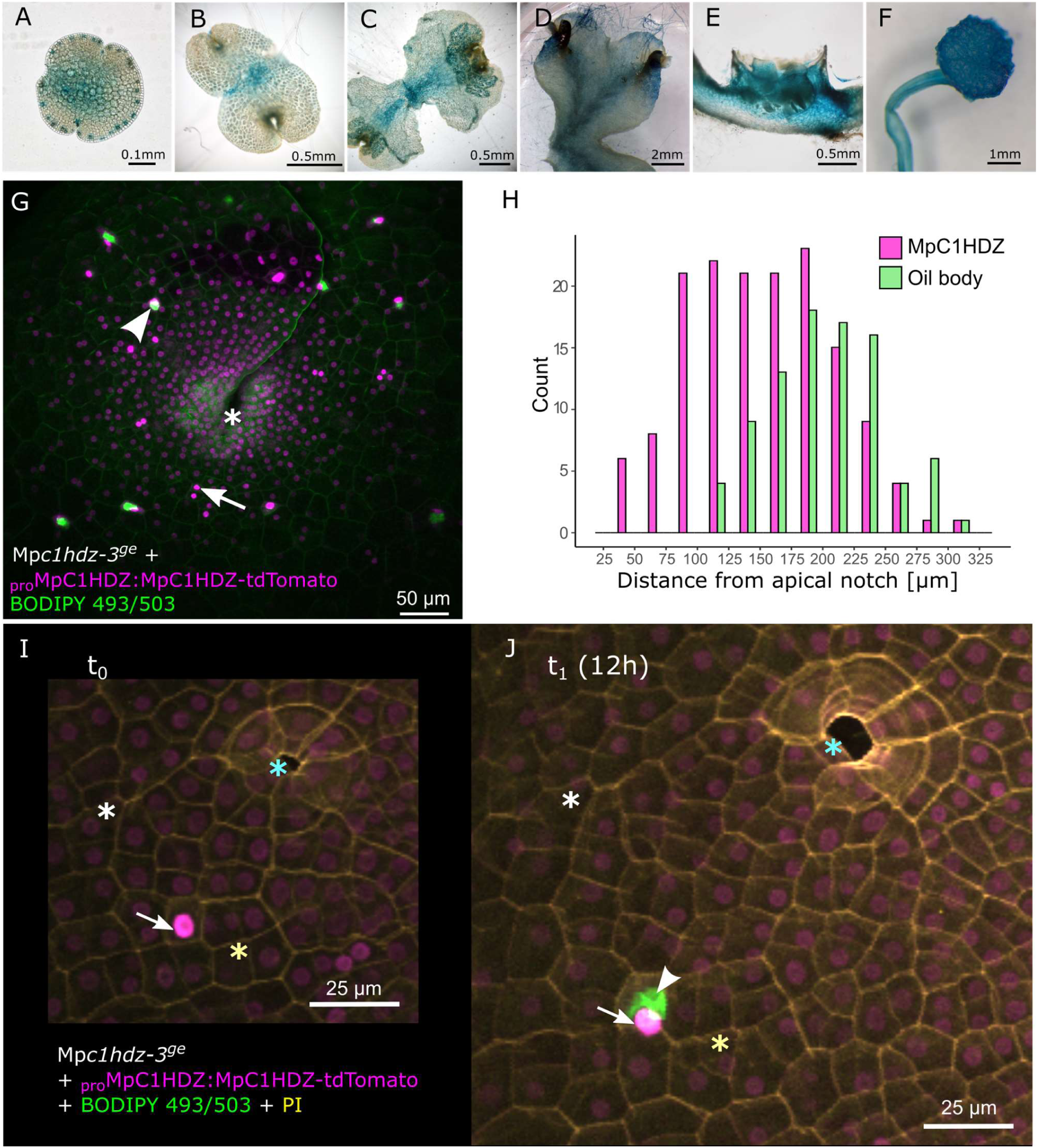
Strong Mp*C1HDZ* expression precedes oil body formation. (A-F) Representative pictures of _*pro*_Mp*C1HDZ*^*4.6kb*^:*GUS* staining (from left to right) of fresh gemmae and 1-week-old plant (Australian ecotype) light microscopy, stereoscopic microscopy of transverse sections of 3-week-old gemma cup grown in non-axenic conditions. (G) A representative apical notch area of Mp*c1hdz-3* gemmaling complemented with _*pro*_Mp*C1HDZ*^*3kb*^:Mp*C1HDZ-tdTomato* and stained with BODIPY 493/503. Sum slices projection of a confocal stack shows _*pro*_Mp*C1HDZ*^*3kb*^:Mp*C1HDZ-tdTomato* signal in magenta and BODIPY in green. Asterisk marks the apical notch, arrowhead marks an oil body, and arrowhead marks an exceptionally bright nucleus. (H) Distribution of MpC1HDZ-tdTomato expression maxima and oil body distances from apical notch in 3 day-old gemmalings, quantified from 2D projections as in G. Quantification method is described in Figure S7. All bright nuclei above a defined MpC1HDZ-tdTomato fluorescence intensity threshold and all oil bodies from 11 replicates were counted. Significance was tested using Wilcoxon rank sum test, *p.* value < 0.05. (I-J) Mp*c1hdz-3* gemmaling complemented with _*pro*_Mp*C1HDZ*^*3kb*^:Mp*C1HDZ-tdTomato* (magenta), stained with PI (yellow) and BODIPY 493/503 (green) and observed using CLSM microscopy in 12 h intervals (Standard deviation z-stack projections). Asterisks of the same color mark the corresponding landmarks in different time-points. Arrow points to nucleus of the cell that developed an oil body (arrowhead) within the next 12 h. All 13 observed transition events are shown in Figure S8 and the event shown here corresponds to number 6 (S8 I, J). Scale bars are shown in the picture.

To further test the role of MpC1HDZ in responses to environmental conditions, we used the transcriptional reporter in a wide range of stressors (Figure S6). Unlike in angiosperms [40], Mp*C1HDZ* was not induced by abiotic stress treatments or ABA (Figure S6). We also tested biotic stress related treatments, such as salicylic acid (SA), wounding, or bacterial elicitors, which did not change the expression activity of Mp*C1HDZ* promoter (Figure S6). However, among environmental conditions known to induce oil bodies, we found a strong induction of Mp*C1HDZ* expression in the meristematic region when plants were grown in non-axenic conditions (soil pots), but not in starvation (Figure S6).

### Mp*C1HDZ* expression maxima precede oil body formation

To understand the dynamics of oil body formation and Mp*C1HDZ* expression, we used a translational reporter _*pro*_Mp*C1HDZ*^*3kb*^:Mp*C1HDZ*-*tdTomato* that can complement the mutant phenotype. As observed in tobacco transient assays, fluorescently labelled MpC1HDZ-tdTomato was also nuclear localized in stable *M. polymorpha* transgenic lines (Figure 2G, Figure S1). In 3 day-old gemmalings, high-expressing cells were located close to the notch in undifferentiated meristematic cells (Figure 2G). MpC1HDZ-tdTomato expression levels dropped gradually from the apical notch to distant cells, except for some isolated cells where MpC1HDZ-tdTomato accumulation became markedly higher compared to neighboring cells (Figure 2G). Using confocal microscopy to quantify the fluorescence of MpC1HDZ-tdTomato and oil bodies stained using BODIPY dye, we found an association between MpC1HDZ-tdTomato accumulation and oil body cells (Figure 2H). Closer to the apical notch sporadic cells strongly expressed MpC1HDZ-tdTomato without BODIPY signal in their vicinity. However, in cells more distant from the apical notch, the brighter MpC1HDZ-tdTomato nuclear signal was associated with BODIPY-stained oil bodies (Figure 2G,H). This initial correlation was further confirmed with cell tracking experiments. We observed that 11/13 newly detectable oil bodies in the field of view formed in cells that had exceptionally high MpC1HDZ-tdTomato expression in previous time point (Figure 2I,J, Figure S8). These results suggest that MpC1HDZ expression maxima precede oil body formation. Taken together, our results indicate that oil body cell differentiation is positively regulated by the MpC1HDZ.

### Mp*c1hdz* mutants show normal abiotic stress responses

Ectopic expression of C1HDZ genes from different angiosperms in Arabidopsis enhances tolerance to water stress such as water deprivation, osmotic stress and salt stress [40, 47-51]. Moreover, some Arabidopsis loss-of-function C1HDZ alleles showed a reduced sensitivity to abscisic acid (ABA) treatments, consistent with a role in water stress tolerance [52, 53].

Using established protocols to manipulate abiotic stress conditions, we applied different treatments including water deprivation, osmotic and salt stress, to quantify the impact on plant growth, *i.e.*: area growth, thallus biomass. Our results showed no altered growth responses of the mutant lines compared to the wild type (Figure 3A-C; Figure S9). We also performed a water-loss experiment of detached thalli and an ABA-dose response germination assay. We did not find a differential response of the mutant in either case (Figure 3D-E). To test whether Mp*C1HDZ* expression is regulated by abiotic stress, we performed RT-qPCR analysis under NaCl and ABA treatments at two time-points, including also other members of the same protein family. Unlike the strong induction observed for Mp*DHN1* [54], Mp*HDZ* expression remained unaltered or were slightly reduced under these experimental conditions (Figure S10).

**Figure 3.**
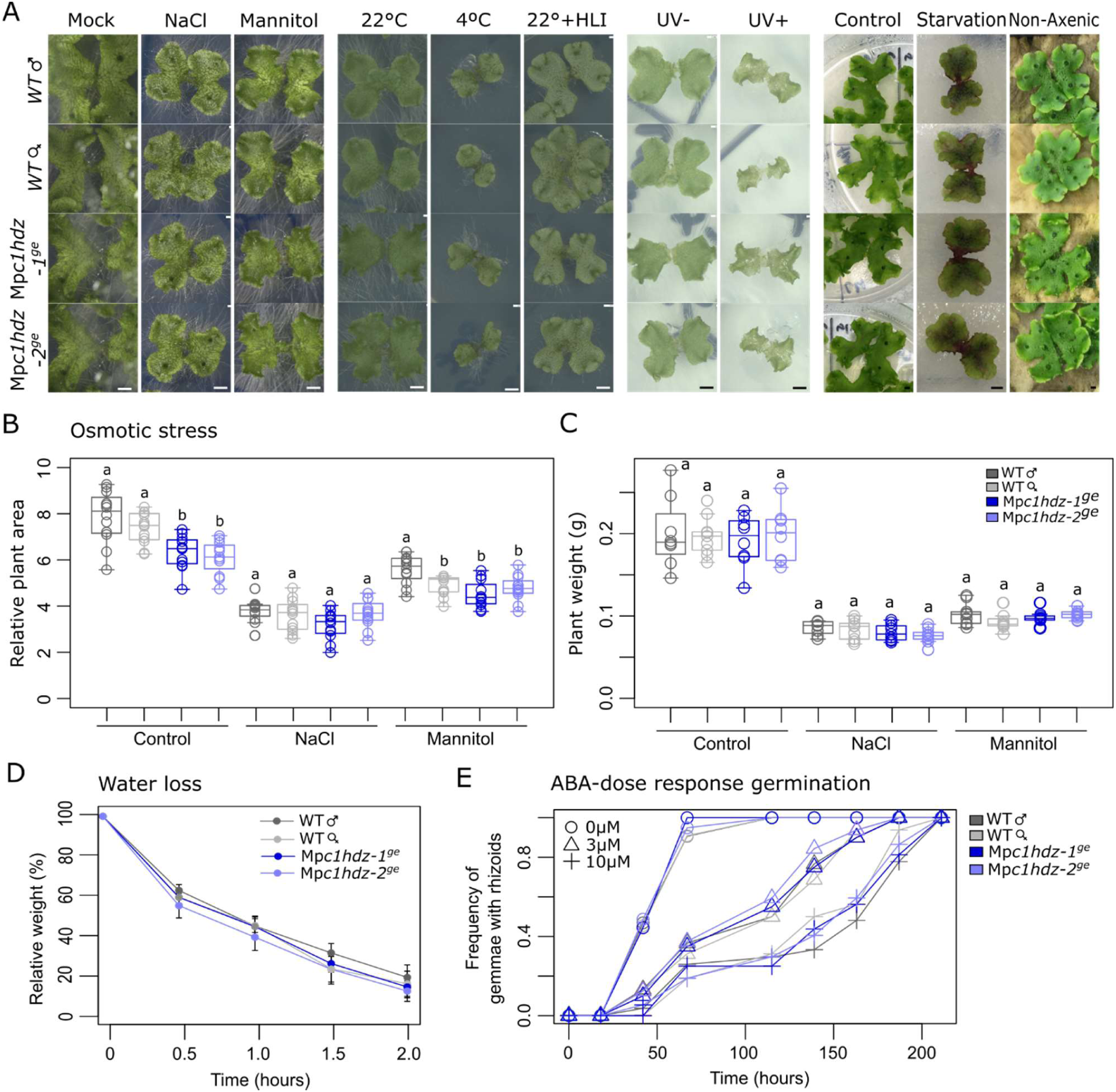
Mp*c1hdz* plants have an intact growth response to abiotic stress. (A) Representative picture of phenotypic effect of abiotic stress treatments on Mp*c1hdz* and wild-type plants as explained in methods section. Treatments are indicated at the top of illustrative pictures. Scale bars (2mm) are indicated in the lower panels. Individual values are indicated as circles. (B-D) Quantification of exposed area (B) and weight (C) of 3-week-old plants under osmotic stress treatments. Individual values are indicated as circles. Full quantifications for each treatment are shown in Figure S7. (D) Water loss experiments were done using 3-week-old wild-type and Mp*c1hdz* plants placed on filter paper towel and weighed every 30 min for 2 h at room temperature. Bars show the average of three biological replicates ± SE of the mean. No statistical difference was observed using ANOVA followed by Tukey HSD after 1 hour of treatment (*p.*value > 0.05). (E) Gemmae dormancy under ABA treatments at different concentrations and time points (left, n ∼ 25). No statistical difference was found testing equal frequencies against wild-type using Exact Binomial Test (*p.*value > 0.05) at 115 h.

Oil bodies have long been proposed to protect liverworts against other abiotic stresses [55-57]. Therefore, we examined the response of Mp*c1hdz* plants to a wide range of other abiotic stressors. We grew plants in axenic control conditions for one week and subsequently transferred plants to different stress environments and measured growth parameters under cold stress, high light irradiance and UV light treatments (Figure S9). We also measured growth parameters in plants subjected to treatments that promote the formation of oil bodies (Figure S9). We did not observe significant differences in growth parameters among wild type and Mp*c1hdz* plants in any of the tested conditions. Overall, our results suggest that neither oil bodies nor Mp*C1HDZ* are involved in the abiotic stress responses of *M. polymorpha* that C1HDZ gene function observed in angiosperms is not conserved throughout land plants.

### Mp*c1hdz* mutants has an impaired biotic defense response

Since oil bodies are rich sesquiterpenes stores and these compounds are known to play a central role in the ecology of plant-biotic interactions, together with old experiments associating the presence of oil bodies to the feeding behavior of snails [26], we investigated if oil bodies play a role in the biotic response of *M. polymorpha*. We performed a choice experiment using wild type and Mp*c1hdz* plants to the terrestrial isopod *Armadillidium vulgare* (pill bug; Crustacea). In this case, we monitored the total thalli area consumed by starved pill bugs for 24 h. The total area lost from mutant alleles was significantly larger compared to respective wild-type plants in both genetic ecotypes (Figure 4A-C, Figure S11). Thus, isopods prefer mutant plants over wild-type, consistent with the possible role of oil bodies as a defense against arthropods herbivory.

**Figure 4.**
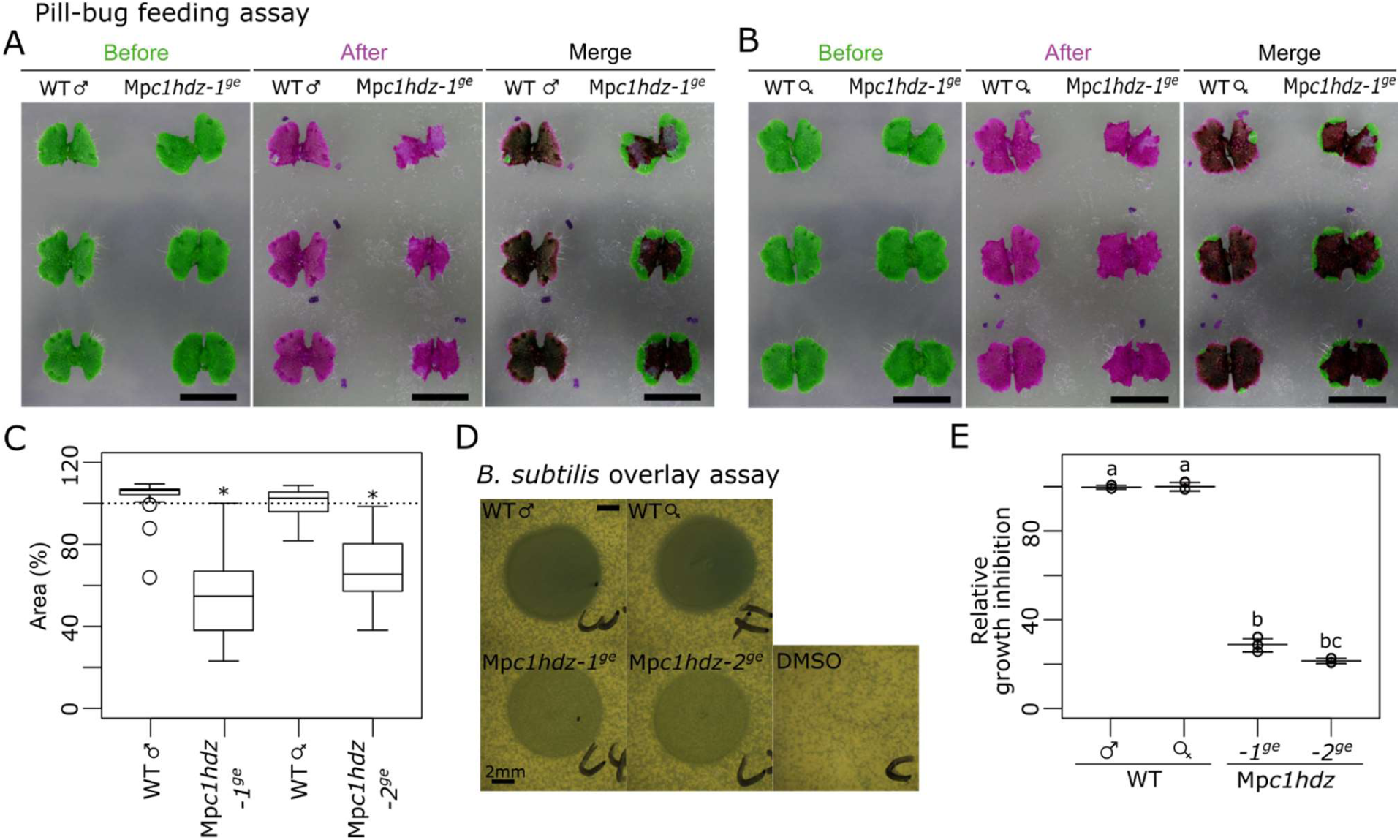
Phenotypic analysis of Mp*c1hdz* plants challenged with biotic stressors. Feeding assay was performed using starved pill bugs (*Armadillidium vulgare*) and 10-day-old thalli were co-cultivated for 24 hours. In order to contrast thallus areas, plants were colored in green (before) and magenta (after). Male wild-type and Mp*c1hdz-1*^*ge*^ thalli (A). Female wild-type and Mp*c1hdz-2*^*ge*^ thalli (B). Scale bars represent 1cm. (C) Thallus damage quantification was performed using ImageJ and data represented as box plots. Thallus area change indicates the ratio of thallus area after and before (n = 36). Statistical differences are marked as asterisks and computed pairwise *t*-test (*p* value < 0.01) comparing each mutant with their respective control. (D) *Bacillus subtilis* overlay assay using ethanolic lipid extracts from wild-type and Mp*c1hdz* mutant thalli dissolved in DMSO. Representative images of experimental results (three replicates were performed with consistent results). Scale bars = 2mm. (E) Quantification of relative growth inhibition as density of colonies relative to wild-type. The graph shows the mean with individual values (n = 3). Statistical difference was tested using ANOVA followed by Tukey HSD *p.*value < 0.01; letters indicate statistically significant groups.

As several compounds produced in oil bodies also have antibiotic activity [32, 58], they may provide further protection against biotic assault. To test this, we applied ethanolic extracts dissolved in DMSO from wild-type and Mp*c1hdz* thalli to the Gram (+) soil bacteria, *Bacillus subtilis*, using an overlay assay. We observed that antibacterial activity was reduced in Mp*c1hdz* extracts in a dose-dependent manner, around one order of magnitude compared to wild-type extracts (Figure 4D,E, Figure S11). Thus, in contrast to abiotic stresses, the striking phenotypic effects observed when challenged with biotic stressors provide convincing evidence that oil bodies are critical for biotic defense responses.

### Mp*C1HDZ* alters the expression of terpenoid-related genes

To better understand the molecular mechanisms underlying the altered defense response of the Mp*c1hdz* mutant, we compared the genome-wide transcript profiles of wild-type and Mp*c1hdz* plants grown in control (axenic) and non-axenic conditions. Here, we show the results of consistently differentially expressed genes between both RNA-seq experiments (Figure 5A,B). In control conditions, we identified 217 down-regulated genes (log_2_(FC) < −1, adj.*p.*value < 0.05) and 131 up-regulated genes (log_2_(FC) > 1, adj. *p* value < 0.05) in Mp*c1hdz* mutant plants (Figure 5, Table S1). Whereas, in non-axenic conditions, we identified 602 genes down-regulated and 267 up-regulated (Figure 5, Table S2). Additionally, we performed enrichment analysis over differentially expressed gene groups using an in-house pipeline of automatically annotated gene lists including GO-terms, biological processes and protein families (Table S3). Among down-regulated genes, there was an enrichment in biological processes associated with secondary metabolism. Moreover, we found enrichment in proteins associated with terpene synthases (TPS, Table S3). Among, up-regulated genes, we did not find any significant enrichment.

**Figure 5.**
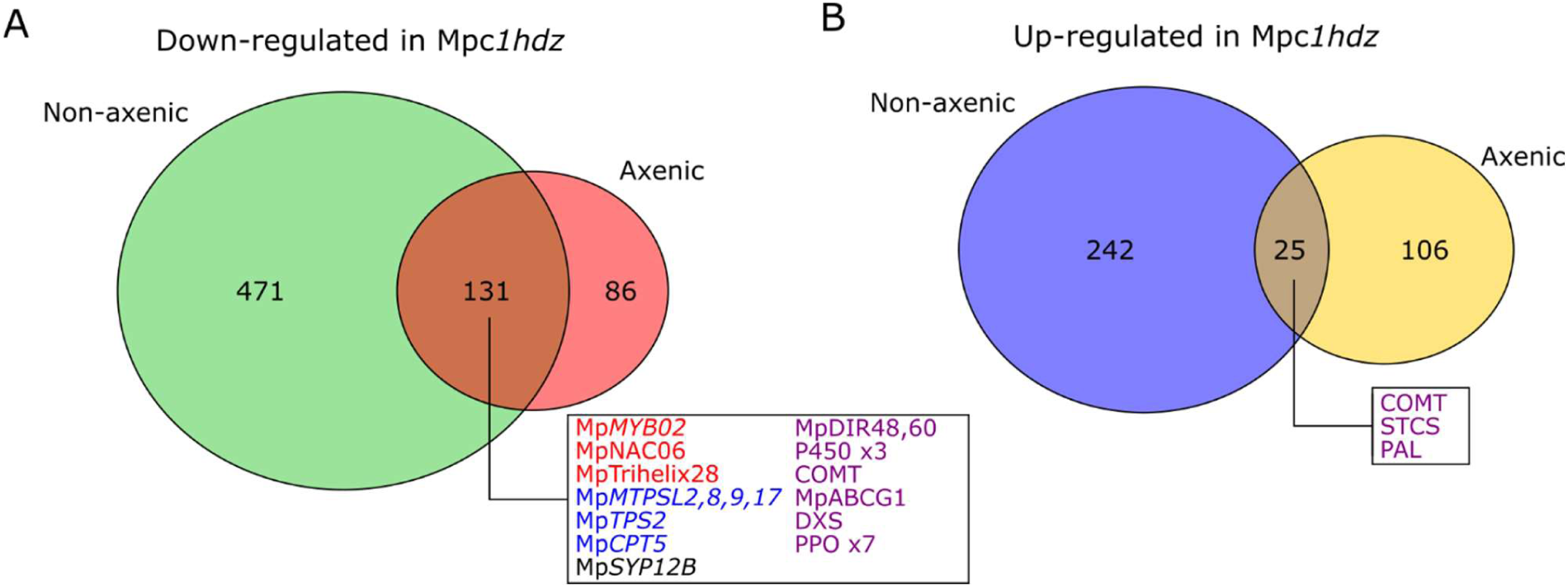
Transcriptomic analysis of Mp*c1hdz* mutants. Venn diagrams of group of genes down-regulated (A, log_2_(FC) < −1, adj.*p.*value < 0.05) and up-regulated (B, log2(FC) > 1, adj.*p.*value < 0.05) in 3-week-old vegetative thalli of Mp*c1hdz* compared to wild type in control (axenic) and non-axenic conditions as described in Methods section. Manually annotated genes associated with TFs (red), terpenoid-related biosynthesis (blue) and other secondary metabolite biosynthetic processes are highlighted. The full data is available at Table S1 and S2.

Within down-regulated genes associated with secondary metabolism (Figure 5A) and down-regulated in both experiments, we found five Mp*TPS* (Figure 5A), including mono- and di-TPS with a bacterial, fungal or plant origin [33, 59]. We also detected other genes involved in the biosynthesis of terpene precursors, including a deoxyxylulose 5-phosphate synthase (*Mp4g14720*, Mp*DXS*) and a cis-prenyltransferase (*Mp3g18510*, Mp*CPT5*) gene (Figure 5A). In addition, three TF genes (*Mp3g07510*, Mp*MYB02; Mp5g01530*, Mp*NAC06* and *Mp2g16340*, Mp*Trihelix28*) are down-regulated (Figure 5A). Mp*MYB02* is known to regulate cyclic bis(bibenzyl) acids biosynthesis [35] which accumulates in oil bodies [15]. The mutant also exhibited significantly reduced expression of Mp*SYP12B* (*Mp4g20670*), a N-ethylmaleimide-sensitive factor attachment protein receptor (SNARE) protein with specific localization at the oil body membrane of *M. polymorpha* [60]. Finally, we also identified other down-regulated genes that might be associated with the secondary metabolism, including a catechol-O-methyltransferase (*Mp5g06850*, Mp*COMT*), two dirigent proteins (Mp3g24140, Mp*DIR48* and Mp3g24210, Mp*DIR60*), an ABC transporter (*Mp8g13070*, Mp*ABCG1*), and cytochrome P450 coding genes.

### Oil body specific terpenoids are depleted in Mp*c1hdz* mutant plants

The strong reduction of oil bodies and the low expression of some *TPS* genes and other genes related to secondary metabolism, led us to explore the terpenoid profile of Mp*c1hdz* vegetative thalli. With the aim of evaluating the impact of environmental conditions on oil body chemistry, we also compared the terpenoid profile of plants grown under control, non-axenic, and starvation conditions (Figure S13A-C). In agreement with previous results [15], GC-FID and GC-MS analyses of thalli hexane extracts showed that non-axenic growth and starvation significantly increased the amount of terpenoid-related compounds compared to wild-type backgrounds, with starvation the most inductive condition (Figure S13D). Interestingly, male backgrounds showed higher levels of terpenoids than female ones, even in the mutant alleles (Table S4).

More importantly, the chemical profile of Mp*c1hdz* thalli revealed a specific decrease of terpenoid-related compounds (**2-7**), such as cis-thujopsene, β-chamigrene, β-himachalene and 5-hydroxy-gurjunene (Figure 6A,B). Most compounds that accumulated under oil body inducing conditions in wild-type plants were concomitantly reduced in mutant plants, suggesting they are located in oil bodies as suggested previously [15, 16] (Figure 6A,B). The most abundant monoterpene d-limonene (**1**) and the unidentified terpenoid U13 (**10**) (Figure 6 C,E; [33]) were strongly depleted in the mutants but they accumulated to similar levels in control and non-axenic conditions (Fig 6C, Figure S14). In contrast, the most abundant diterpene, phytol (**11**) [33] (Figure 6D), fatty acid compounds (**8**,**9**,**12**,**13**) (Figure 6E) and phytosterols, also present in plant extracts, did not significantly change in the mutant or among growth conditions (Table S4). Taken together, these results support that oil body-localized terpenoid biosynthesis is affected in Mp*c1hdz* mutants.

**Figure 6.**
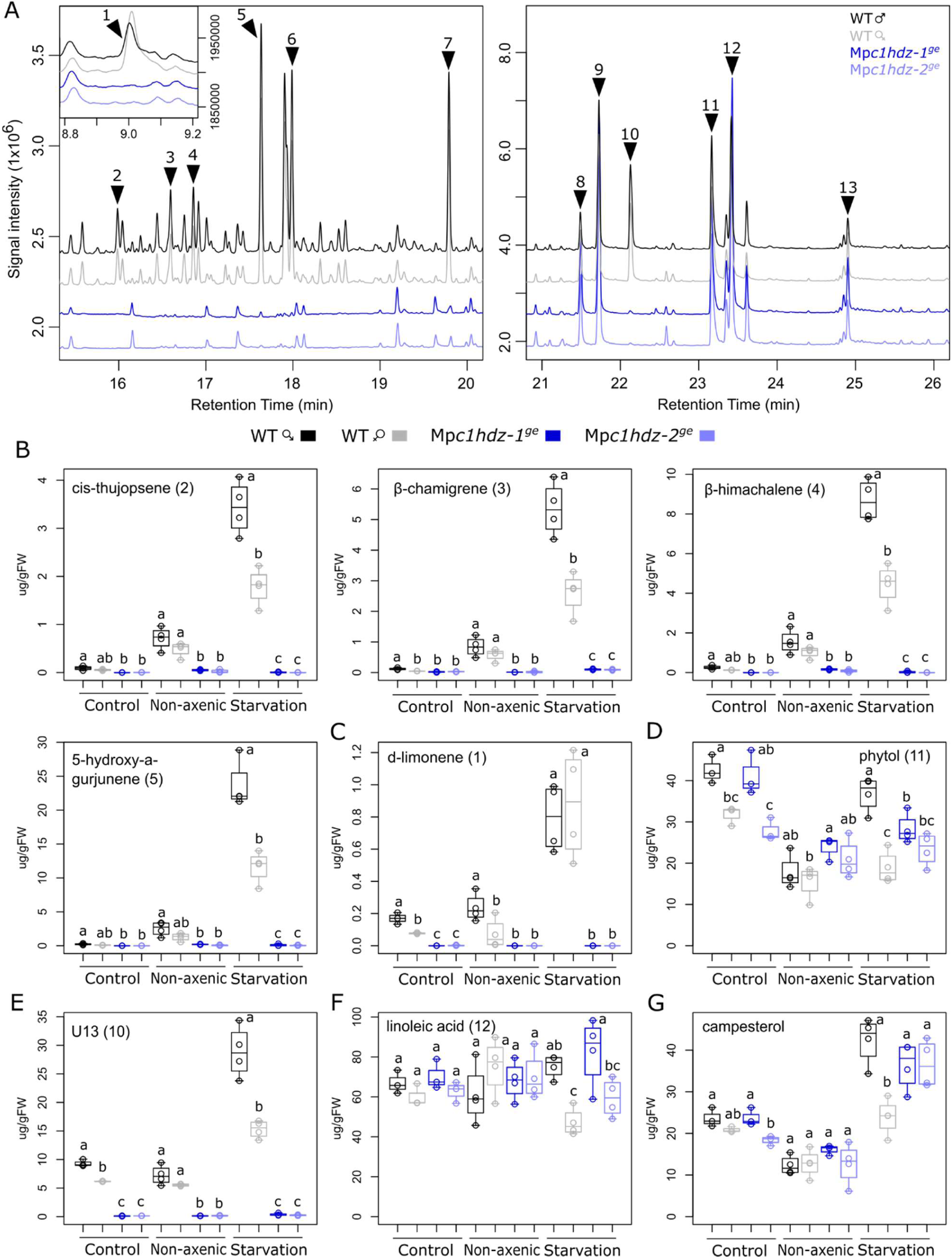
GC-FID analysis of Mp*c1hdz* mutants. (A) Representative GC-FID chromatogram obtained from hexane extracts under starvation treatment of Mp*c1hdz* and wild-type plants. Three different ranges are shown. Arrow points represents metabolites: limonene (**1**), cis-thujopsene (**2**), β-chamigrene (**3**), β-himachalene (**4**), 5-hydroxy-gurjunene (**5**), viridiflorol (**6**), cyclocolorenone (**7**), stearic acid methyl ester (**8**), palmitic acid (**9**), unidentified terpenoid U13 (**10**), phytol (**11**), linoleic acid (**12**), timnodonic acid (**13**). Quantification of representative major metabolites in Mp*c1hdz* and wild-type plants under three different treatment as described before (control, non-axenic, and starvation) and represented as boxplots (n = 3-4): total sesquiterpenes (B), monoterpenes (C), diterpenes (D) and fatty acids (E). Statistical difference among genotypes was tested using ANOVA followed by Tukey HSD *p.*value < 0.05 for each treatment; letters indicate statistically significant groups.

## DISCUSSION

C1HDZ genes have been previously characterized for their roles during abiotic stress response in angiosperms, more specifically to water stress tolerance in Arabidopsis, coffee, cotton, rice and sunflower [40]. Likely due to functional redundancy in angiosperms, the majority of C1HDZ mutants in angiosperms exhibit subtle phenotypes [39], with some exceptions. Homologs of *LATE MERISTEM IDENTITY 1/REDUCED COMPLEXITY* alter leaf morphology via local growth repression at early developmental stages [61, 62]. Given that *M. polymorpha* possesses a single *C1HDZ* gene, this is the first time that a full loss-of-function mutant is generated for this subfamily. Interestingly, the lack of a functional MpC1HDZ did not generate major pleiotropic effects as observed for other TF families in *M. polymorpha*. Using a wide array of experimental set-ups, we found no evidence supporting a physiological role for Mp*C1HDZ* in the abiotic stress response of *M. polymorpha* (Figure 3, Figure S9). Further, in the moss *Physcomitrella patens*, only 1 out of 17 C1HDZ genes responded to dehydration and mannitol treatments and the only studied Pp*c1hdz* mutant presented a divergent function [45, 63]. These data suggest that the physiological role of C1HDZ genes in response to abiotic stress may be largely a vascular plant phenomenon.

### Mp*C1HDZ-*mediated depletion of oil bodies affects arthropod herbivory

We found compelling evidence that Mp*C1HDZ* modulates, directly or indirectly, the differentiation of oil body cells. Consistently, loss-of-function Mp*c1hdz* alleles in two ecotypes exhibited a strong reduction in the number of oil bodies (Figure 1). Early experiments showed that the number and size of oil bodies in liverworts remained mostly unaffected in a variety of environmental conditions, such as light, temperature and limiting N or P [8, 64, 65], and that once formed they can be stable in herbarium specimens [10]. It was recently shown, however, that axenic growth resulted in a significant reduction in oil body cell formation, with a substantial induction upon a shift to non-axenic growth, such as cultivation on soil substrate where liverworts are challenged with biotic stressors [15]. We found that Mp*c1hdz* plants exhibit a severe reduction of oil body formation under all tested environmental conditions. However, the presence of a limited number of oil bodies in the mutants suggests that other transcriptional regulators could be relevant along with MpC1HDZ.

We explored several environmental conditions where oil bodies could play a physiological role. Whereas we found no evidence that Mp*c1hdz* plants were compromised in the response to an array of abiotic stressors, mutant plants were more susceptible to herbivory by the isopod *A. vulgare* (Figure 4C) and mutant plant extracts were less efficient controlling the growth of *B. subtilis* (Figure 4D). Thus, our results provide genetic evidence supporting a role for oil bodies in liverwort defenses against arthropod herbivores, supporting the hypothesis formulated more than a century ago [26].

### What is the ancestral function of oil bodies?

Oil bodies are a synapomorphy of liverworts, and thus arose in the ancestral liverwort [7], likely placing their origin as early as the Ordovician and raising the question of their ancestral function. Whereas in liverworts where all cells contain oil bodies an antimicrobial role might be plausible, in liverworts with specialized oil body cells (idioblasts) a role in herbivory defense appears more likely. However, the ancestral condition in liverworts, scattered or ubiquitous, is equivocal. There is evidence from the fossil record that the selection pressure of herbivory was acting on ancestral liverworts. Middle Devonian liverwort fossils showing undigested tissues rich in opaque cells resembling the oil bodies of extant liverworts provides indirect evidence of a role in deterring herbivory [27]. Whereas an earlier liverwort fossil record is missing, evidence of arthropod feeding on spores and stems of late Silurian-early Devonian plants has been reported [66]. Furthermore, evidence of herbivory corresponds closely with fossil evidence of plant organ evolution, implying either an immediate herbivory response or an extension of earlier plant-herbivore interactions. Thus, the anti-herbivory function of terpenoids and other secondary metabolites concentrated in oil bodies may date to the ancestral liverwort as was also recently suggested [67]. More broadly, as these compounds also play a central role in the ecology and evolution of the biotic response of seed plants [68], the origin and diversification of these secondary metabolites might have facilitated the evolution of early land plants via modulation of their interactions with the biotic environment.

### Biochemical properties of Mp*c1hdz* mutants

As expected, the paucity of oil bodies in Mp*c1hdz* plants resulted in a marked reduction in specific terpene-related compounds (Figure 6). It has been demonstrated that the sesquiterpenoids and bis(bibenzyls) are accumulated in the oil bodies [15]. These compounds present cytotoxic activity against bacteria and fungi, mammalian and plant cells [14, 15, 29-31] but also have other medical applications [58]. Indeed, such bioactivity is the likely reason *M. polymorpha* is mentioned at least as early as a Greek herbal in the first century as a surface poultice to prevent inflammation of wounds [69]. *In planta*, while we provide evidence of a role for oil bodies against isopods, additional functions in fungal or bacterial resistance have not been ruled out. However, that liverwort species with oil bodies in every cell form mycorrhizal fungal interactions [70] indicates fungi can avoid their cytotoxicity, perhaps due to oil bodies being membrane bound organelles.

### Oil body cell differentiation depends on MpC1HDZ

In *M. polymorpha* thalli, idioblasts emerge close to the apical notch in a proportion that later decreases due to unequal cell division rates [8]. Nonetheless, the precise mechanism of oil body differentiation and formation is still unclear. Here, we described Mp*c1hdz* mutants that not only have a severe reduction of oil bodies but also do not differentiate idioblast cells (Figure 1). Transcriptional analysis of Mp*c1hdz* thalli demonstrated that several down-regulated genes encode proteins located in the oil body (TPS, MTPSL and MpSYP12B; [16, 60]), together with other secondary metabolism related genes that could be directly activated by MpC1HDZ (Figure 5). It is unclear whether the lethality encountered in Mp*C1HDZ* gain-of-function mutants results from ectopic production of cytotoxic compounds, ectopic oil body cells, or another defect.

As oil bodies form at low frequency in Mp*c1hdz* mutants, other transcriptional regulators could be important for oil body cell differentiation. The R2R3-MYB TFs Mp*MYB02* and Mp*MYB14* control the bis(bibenzyl) and auronidin biosynthesis, respectively [35, 37]. Mp*c1hdz* mutants showed a strong repression of Mp*MYB02* expression, whereas Mp*MYB14*, important for the defense against the oomycete pathogen *Phytophthora palmivora* [71], remained unchanged (Table S1, S2). That Mp*C1HDZ* expression is not altered in Mp*MYB02* gain-of-function alleles [35] suggests that Mp*MYB02* is downstream MpC1HDZ activity, however, it is unknown whether MpMYB02 is also specifically expressed in oil body cells or participates in oil body cell differentiation and terpenoid biosynthesis. In contrast, we did not find any relationship of Mp*C1HDZ* with Mp*MYCX/Y*, which is associated with the defense-related induction of sesquiterpenoid biosynthesis in response to herbivory [67], or with ABA signaling, which is associated with bis(bibenzyl) increase under UV-C light stress [38].

Based on our observations, we propose a model for oil body cell differentiation (Figure 7). Undifferentiated cells derived from the notch express and accumulate MpC1HDZ to high levels; stage 1) a higher expression level of MpC1HDZ in one of the cells triggers differentiation of an oil body cell, while MpC1HDZ accumulation in neighboring cells decreases; stage 2) oil body forms in cells with high MpC1HDZ expression and oil body components are biosynthesized; stage3) oil bodies mature while MpC1HDZ expression fades, and growth displaces them from the notch as non-idioblast cells continue dividing (Figure 7). We propose that MpC1HDZ mediates the developmental program of oil body cell differentiation. Notably, our results do not exclude that other transcription factors may act earlier than MpC1HDZ in oil body cell differentiation. Investigating how integration of different transcriptional and environmental inputs promotes oil body cell development and differentiation will help with further understanding of this process. Such information will also pave the way for manipulating oil body numbers and composition for biotechnological purposes.

**Figure 7.**
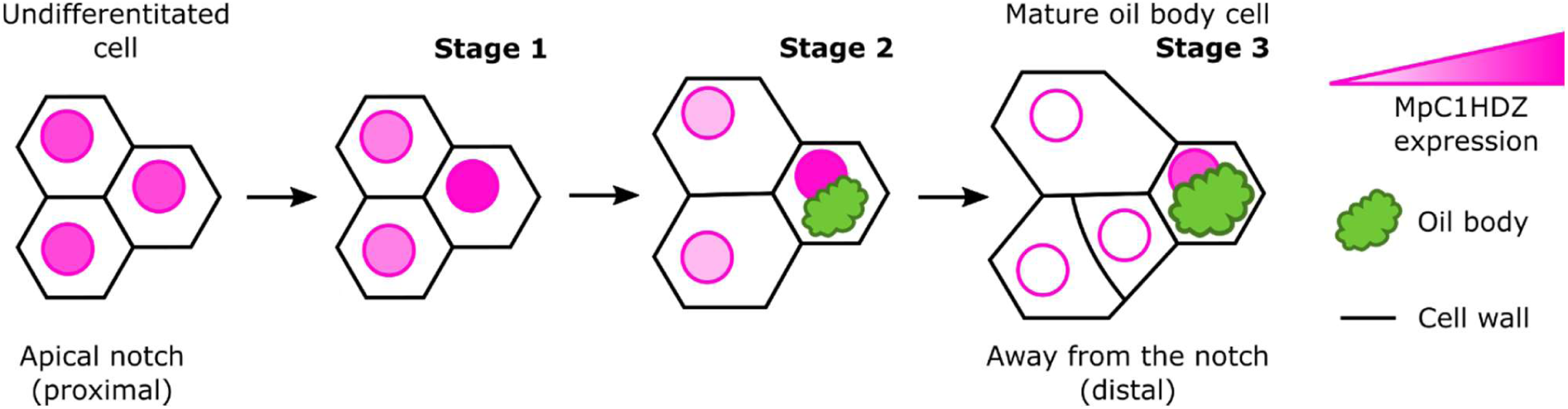
Proposed model for MpC1HDZ dynamics of oil body cell differentiation. All cells in proximity of an apical notch express MpC1HDZ. While most of the cells gradually loose MpC1HDZ expression, some cells start expressing MpC1HDZ stronger than their neighbors (stage 1). These cells can form an oil body (stage 2). With distance from apical notch and maturation of oil body cell, MpC1HDZ expression fades (stage 3). During oil body cell development, neighboring cells continue expanding and dividing, while oil body cells remain smaller.

## MATERIAL AND METHODS

### Plant material and growth conditions

*Marchantia polymorpha* sp. ruderalis Australian (Melbourne) ecotype [72] and BoGa ecotype (obtained from Botanical Garden in Osnabrück, Germany) [73] were cultivated from gemmae under axenic conditions on vented 9 by 2.5 cm Petri dishes containing ½ strength Gamborg’s B5 medium pH 5.5 solidified with 1 % (w/v) agar, referred as normal media. Australian ecotype was grown in growth chambers with 16/8 h photoperiod, white light at 80-100 µmol m^-2^s^-1^ at 22°C. BoGa ecotype was grown in growth chambers with continuous white light at 50 µmol m^-2^ s^-1^ at 22°C. Sexual reproduction was induced using white light supplemented with far-red light (730 nm, Cree Inc. LEDs) as described before [74]. Plants were crossed in sterile conditions using autoclaved water for spermatozoid transfer. Crossing was repeated twice and plants were returned to long day condition until sporangia were ready for harvest. Sporangia were dried and stored at −80°C.

### Marchantia transformation

*M. polymorpha* spore transformation was performed as described previously using both ecotypes [75]. Briefly, *M. polymorpha* spores were germinated and grown in 25 mL ½ strength liquid Gamborg’s B5 with 2%(w/v) sucrose, 0.1%(w/v) Casamino acids and 0.03%(w/v) L-Glutamine for 10 days prior to co-cultivation with 1 mL of an *Agrobacterium tumefaciens* (GV3001) suspension, harboring binary plasmids, along with acetosyringone to a final concentration of 100 µM. Sporelings were plated on selection media (10 µg/mL Hygromycin B with 200 mg/mL Timentin). After two weeks, T1 lines were transferred for one more week to new selection plates before being transferred to standard growth plates for gemmae production. Gemmae (G1 or G2 generation) from several independent primary transformants (T1 generation) were analyzed for the presence of the transgene by PCR and sequencing, with some selected for further analysis.

Translational reporters were transformed into Mp*c1hdz-3*^*ge*^ mutant using *Agrobacterium*-mediated transformation of regenerating thalli [76]. Briefly, plants were grown from gemmae on standard medium in continuous light for 14 days. 14 day-old thalli were cut into four pieces without apical notch and cultured for 3 days on standard medium with 1 % sucrose to induce regeneration. Regenerating thalli were co-cultured with *A. tumefaciens* GV3101 in liquid medium with 100 μM acetosyringone with agitation. After three days the plants were washed six times with autoclaved water, incubated for 30 min in 1 g/L cefotaxime and plated on selective plates containing 0.6 μM chlorsulfuron and 120 μg/mL cefotaxime. After transformants appeared, they were transferred to new selective plates. G2 generation was analyzed for fluorescence and presence of the transgene by PCR.

### Genetic constructs

All primer sequences are available in Table S5.

### gRNA design and cloning

Synthetic guide RNAs (gRNA) targeting genes of interest were co-expressed on independent T-DNAs using pGE010 plasmid [77, 78]. Design of specific gRNAswas performed such that putative off-targets (*M. polymorpha* genome assembly v3.1) lacked an associated protospacer adjacent motif (PAM) site. For single gRNAs transformation, primers MpC1HDZ_gRNA1 were cloned downstream of the MpU6 promoter, excluding PAM sites, in pMpGE_En03 using BsaI sites and were co-expressed with Cas9 [77]. For double gRNAs transformations, gRNAs were designed using the online tool http://marchantia.info/casfinder. The first gRNAs (gRNA-wg-1 or gRNA-HD-1) were cloned into pMpGE_En04 using BsaI sites [79]. Second gRNAs (gRNA-wg-2 or gRNA-HD-2) were cloned into pMpBC_GE14 using BsaI sites. Both vectors were digested using BglI. The pMpGE_En04 backbone with the first gRNA and the fragment with the second gRNA were ligated, resulting in pMpGE_En04 with both gRNAs in tandem each under the control of the MpU6 promoter. These vectors were used in a Gateway LR reaction with the binary vector pMpGE010 featuring Cas9 under the control of the EF1 promoter [77-79].

### Mp*C1HDZ* cloning and genetic constructs

The Mp*C1HDZ* (*Mp3g02320*) coding sequence from *M. polymorpha* Tak-1 genome v3.1 (marchantia.info) was used to design specific primers. The cDNA was subcloned into pENTR-DTOPO (Thermo Fisher, Waltham, MA, US; pENTR-MpC1HDZ) using primers MpC1HDZ_CACC_fw and MpC1HDZ_rv. The CDS was mutagenized using primers MpC1HDZ_gibson_fw and MpC1HDZ_mut_rv, and MpC1HDZ_mut_fw and MpC1HDZ_gibson_rv in order to make it resistant to the gRNA. Fragments were purified and used as template for a subsequent amplification using primers MpC1HDZ_gibson_fw and MpC1HDZ_gibson_rv and cloned in a pENTR2B (Thermo Fisher, Waltham, MA, US) using NEBuilder HiFi DNA Assembly Master Mix (New England BioLabs Inc., UK, pE2B-MpC1HDZ_mut_). Later, it was subsequently recombined using the Gateway LR Clonase II Enzyme Mix (Thermo Fisher, Waltham, MA, US) according to the manufacturer’s instructions into different destination vectors: pMpGWB403 (_pro_EF1:MpC1HDZ_mut_), pMpGWB406 (_pro_35S:MpC1HDZmut-Citrine) [80]. As a control, a 4.6 kb promoter region of MpC1HDZ was amplified with primers proMpC1HDZ_gibson_fw and proMpC1HDZ_gibson_fw as described below. Both fragments were used for cloning in a pENTR2B with NEBuilder HiFi DNA Assembly Master Mix and recombined in pGWB401 (_*pro*_Mp*C1HDZ*^*4.6kb*^-Mp*C1HDZ*_*mut*_).

For overexpression in tobacco, pENTR-MpC1HDZ was recombined in the pFK247 expression vector derived from the pGreen vector series [81] for transient expression in tobacco plants.

### Transcriptional reporters

The promoter sequence of Mp*C1HDZ* was obtained from Phytozome. A 4.6 kb fragment of the promoter was amplified using primers proMp*C1HDZ*-NdeI-fw and proMp*C1HDZ*-SalI-rv and cloned into pCRII-TOPO. It was subsequently subcloned into pRITA, which has a multiple cloning site upstream of a GUS/UidA reporter gene and *nos* terminator [82], using NdeI and KpnI. Finally, _*pro*_Mp*C1HDZ*^*4.6kb*^:*GUS* was cloned into pSKF-KART using NotI [72].

Alternatively, a 3 kb sequence upstream of Mp*C1HDZ* start was amplified from BoGa DNA using primers with attB1 and attB2 sites. The subsequent PCR product was recombined with pDONR201 in BP reaction. The entry clone was recombined in an LR reaction with pMpGWB104 to obtain GUS reporter [80].

### Translational reporters

To obtain translational reporters, cDNA was obtained from total RNA extracted from BoGa ecotype thalli. Mp*C1HDZ* was amplified from cDNA using a specific forward primer and polyT reverse primer (Roche) and ligated into pGEMT. The amplified CDS starts at position 3 and ends at position 720 relative to the genome prediction and therefore likely represents a shorter transcript. Mp*C1HDZ* CDS was amplified again from this plasmid to add *SpeI* sites. The 3 kb sequence upstream of Mp*C1HDZ* CDS start was amplified from BoGa DNA and ligated into pGEMT vector. Mp*C1HDZ* CDS was inserted behind the 3 kb MpC1HDZ promoter using SpeI restriction sites. The 3 kb Mp*C1HDZ* promoter and Mp*C1HDZ* CDS combination were amplified using primers with attB1 and attB2 sites and recombined in BP reaction with pDONR201. The entry clone was recombined in LR reaction with pMpGWB307 destination vector to obtain citrine fusion, or with pMpGWB329 to obtain tdTomato fusion [80].

### Histology and microscopy

Fresh *M. polymorpha* gemmae were resuspended in 0.1% (v/v) Triton-X100 and observed using a compound microscope (Nikon Eclipse E200, Nikon Instruments Inc., Melville, NY, USA; or Zeiss Axio Imager M2) or Leica MZ10F stereomicroscope (Leica Microsystems, Wetzlar, Germany; or Nikon SMZ1500). In the case of thallus and cup cross sections, fresh thalli were sliced into 0.5-0.8 mm thick slices manually using a razor blade and observed in the compound microscope.

Oil body staining was performed using a 10 mg mL^-1^ Nile Red dissolved in phosphate buffered saline (PBS) just before use as described before [15]. This solution was vacuum infiltrated for 1 min, incubated 10 min at room temperature and then rinsed with 0.1% (w/v) Triton-X100 solution. For BODIPY staining, the stock solution was diluted 1000 times in PBS to prepare the working solution. Plants were covered with BODIPY for 10 min in darkness and washed three times with PBS.

The samples were observed using GFP filter under fluorescent microscope (Zeiss Axio Imager M2). Oil bodies in gemmae were manually counted from images. Thallus area was quantified at different time intervals from pictures with ImageJ software. Thallus growth rate is the proportion between final and initial area of the thallus. All data was analyzed and plotted in R with built-in packages.

### GUS staining

GUS assays were performed as described previously [72]. Plants were immersed in GUS staining solution (100 mM Potassium Ferrocyanide, 100 mM Potassium Ferricyanide and 1 mM X-Gluc) and after applying 2 rounds of vacuum for 1 min, the plants were incubated for 1.5 h to 2 h at 37°C in darkness, cleared with ethanol, and imaged using a compound microscope or stereoscopic microscope. BoGa ecotypes were stained similarly but with slightly different GUS staining solution composition (50 mM phosphate buffer, 10 mM EDTA, 50 mM Potassium Ferrocyanide, 50 mM Potassium Ferricyanide and 1.4 mM X-Gluc) and the stain was vacuum-infiltrated for 10 min before incubation at 37 °C.

### Oil body cell and oil body distance measurement

For live imaging, gemmalings were grown on standard medium in small Petri dishes (6 cm diameter). 3 day-old gemmalings were stained with BODIPY 493/503 working solution (dissolved in water) for 3 min. Approximately 30 µL dye was used to cover a gemmaling. Samples were rinsed with water two times, covered with water and observed using Leica TCS SP8 confocal microscope and 25x 0.95 NA water dip-in objective. To observe fluorescence, 561 nm excitation and 570 - 600 nm emission was used for tdTomato and 488 nm excitation and 503 – 533 nm emission was used for BODIPY in sequential scan mode. All samples were imaged using the same settings. Images were analyzed in Fiji software. First, all stacks were trimmed to the same size relative to the estimated position of the apical cell along the z-axis. Prior to applying “sum slices” projection to the trimmed stacks, black and white pixels were added in a lower left corner of each slice as a calibration for lowest and highest possible values. In these projections, some nuclei visually stood out as much brighter than the surrounding nuclei. To visualize only these nuclei, all projections were processed as follows: format was set to 16 bit, intensities were multiplied by 100 and smoothed once. For tdTomato channel, brightness & contrast values were set to show only the pixels with intensity value of 10 000 or more. To assess the distances of bright nuclei and oil bodies from the apical notch, distance bins were created using Concentric circles plug-in. Circles were centered in the middle of the apical notch of each image and placed in 25 μm distance intervals. Oil bodies and bright nuclei were quantified for each of the distance bins. Data was visualized and statistically analyzed in R. To visualize apical notch surface in 3D, a surface mesh was created in MorphoGraphX program using signal from BODIPY channel [83].

### Oil body cell tracking

For live imaging, gemmalings were grown on normal media in small Petri dishes (6 cm diameter). A few droplets of water were added around the rim of the medium 30 min before imaging, to soak the agar and prevent sample movements during the imaging. Gemmalings were stained with BODIPY 493/503 working solution (dissolved in water) for 5 min. Approximately 30 µL dye was used to cover a 5-day-old gemmaling. 15 µL of 0.1 % propidium iodide (PI) was added directly to BODIPY and removed after 30 s. Samples were rinsed with water three times, covered with water and observed using Leica TCS SP8 confocal microscope and 25x 0.95 NA water dip-in objective. To observe fluorescence, 488 nm excitation and 503 – 523 nm emission was used for BODIPY, 552 nm excitation and 560 – 580 nm emission was used for tdTomato, and 552 nm excitation and 620 – 640 nm emission was used for PI. The two lasers were used in sequential scan (between stacks), and Lightning mode was used for acquisition. Standard deviation projections of z-stacks were generated in Fiji and used for oil body cell tracking. In case of doubt about the identity or lineage of an object, the stacks were observed in MorphoGraphX.

### Plant stress treatments

Wild-type and Mp*c1hdz* mutant gemmae were grown in agar plates containing normal media. After 1-week growth in control conditions, each plant was transferred with tweezers to a new plate or pot to be exposed to different treatments. Plant areas were recorded before and after the treatment to estimate the impact on the relative growth rate of each genotype. Plant growth rate is the proportion between final and initial area of the thallus. Pictures of whole plants were taken at different time intervals using a Leica MZ10F stereomicroscope. Thallus area was quantified from pictures with ImageJ software. For cold treatments and high light intensity (HLI) plants were transferred to plates containing the normal media (3 plates with 12 plants for treatment and 3 plates as control). Control plates remained in control conditions. Cold treatment was performed at 4ºC in a chamber with similar light and photoperiod conditions and observed after 1 week. HLI treatment was performed in the same chamber but 1000 µmol m^-2^ s^-1^ LED lights with the same photoperiod and observed after 1 week. For UV treatment plants were transferred to normal media and exposed to a laminar flow bench UV-C light for 1 h with the lid open (UV+) or closed (UV-), after that they were recovered in control conditions and observed after 1-week. For osmotic stress, normal media was supplemented with solutions of NaCl or mannitol at a 50mM final concentration or water (mock). Plants were grown in control conditions for 18 days. For starvation, plants were transferred to 1/100 Gamborg’s B5 media (1% (w/v) agar, pH 5.5), grown in control conditions, and observed after 2 weeks (normal media was used as control). For non-axenic growth, plants were transplanted to pots with rockwool as substrate in a growth chamber in similar temperature and photoperiod as control conditions with irrigation.

The water loss experiment was performed as in Re *et al.* [84]. Briefly, 15-day-old wild-type and Mp*c1hdz* plants grown in control conditions and normal media were detached from the agar, placed in a paper towel, and weighed every 30 min during 2 h at room temperature. The percentage (%) of water loss was calculated as follows: [initial weight – weight]/ [initial weight] × 100.

### Pill bug feeding assay

Feeding assay was performed according to Nakazaki *et al*., with slight modification [85]. Gemmae were cultured on 1/2 Gamborg’s B5 medium (Nihon Pharmaceutical, Tokyo, Japan, Cat#399-00621) containing 1%(w/v) agar (Nacalai tesque, Kyoto, Japan, Cat#01028-85) and 1%(w/v) sucrose (Wako Pure Chemical, Osaka, Japan, Cat#196-00015) for 5 days at 22°C under continuous white fluorescent light (50 µmol m^-2^s^-1^). Five-day-old thalli were transferred onto 1/2× Gamborg’s B5 medium containing 1%(w/v) agar without sucrose and cultured for additional 5 days under the same condition above, or 8 days in case of BoGa ecotype. Pill bugs (*Armadillidium vulgare*) were collected at the Myodaiji area of National Institute for Basic Biology (Aichi, Japan). Before the assay, pill bugs were maintained on Prowipe (Daio Paper, Tokyo, Japan) moistened with sterilized water for 48 h at 22ºC under dark condition to be starved. Six pill bugs were introduced into each medium plate containing the 10-day-old thalli, or 13-day-old thalli in case of BoGa ecotype, and kept together for 24 h under the dark condition at 22ºC. The thallus areas of *M. polymorpha* were calculated using ImageJ (National Institute of Health, https://imagej.nih.gov/ij/), and figures were processed with Photoshop software (Adobe systems, USA).

### Ethanol extraction and overlay assay

Wild-type and Mp*c1hdz* gemmae were grown in agar plates containing normal media and control then transferred to pots using soil and vermiculite (1:1 parts) and grown in control conditions with irrigation for one month. Plant materials were snap-frozen with liquid N_2_, and ground using a mortar and pestle (∼5 g fresh weight). The plant powder was extracted with EtOH for 24 h in agitation. Debris was removed by filtration and removal of solvent was performed using a SpeedVac. The stock solution of crude extracts was prepared as 10 mg mL^-1^ in DMSO. *Bacillus subtilis* overlay assay was performed as described before [86]. For quantification purposes, pictures of bacterial growth inhibition were taken using stereoscopic microscope (Leica MZ10F) and calculated as density of colonies.

### RNA-seq

Total RNA isolated from 3-week-old thalli (QIAGEN RNAeasy Plant Kit 5) from each genotype was assessed for quality with a Nanodrop spectrophotometer and a Bioanalyzer 2100 microfluidics system (Agilent 6). For expression analysis in control conditions, one sample per genotype was used (Mp*c1hdz-1*^*ge*^, Mp*c1hdz-2*^*ge*^, WT female, WT male) and mutant and wild-type were treated as replicates. For expression in non-axenic conditions, plants were grown as described before and three biological replicates per genotype were taken from Mp*c1hdz-2*^*ge*^ and WT female. Library preparation substrates were enriched for mRNA (Illumina TruSeq Stranded mRNA technology). Sequencing used the Illumina NextSeq500 7 in HighOutput mode (20 million single-ended 75 b reads per sample). Reads were mapped onto the Marchantia v3.1 (http://marchantia.info/) assembly using TopHat v2.1.0 for Galaxy [87]. Resulting BAM files were loaded onto IGV 1.3.1 to produce Sashimi Plots [88]. A count matrix of average raw reads/gene/sample was created using HTseq-code using Galaxy [89]. Raw reads were normalized by total million reads per library (RPM) and transcript lengths in kb (RPKM). The edgeR package was used for differential gene expression analysis using Galaxy (Table S1 and S2) [90]. Summary statistics of RNA-seq analysis are available in Table S6.

For GO term and protein family enrichment analysis, the algorithm and annotations were as described previously [91]. Gene names from Marchantia v5.1 [92] was assigned in accordance to gene corresponding table available at http://marchantia.info/nomenclature/ [2]. Genes were annotated using information nomenclature previously published [93] or available at MarpolBase.

### Hexane extraction and gas chromatography

Samples of WT and transgenic plants (3-4 samples per genotype) were collected from 3-week-old plants in different treatments and snap frozen in liquid nitrogen. Each sample was ground with a pestle in a mortar containing liquid nitrogen. 300 µL of n-hexane was added to ∼400 mg of ground tissue, centrifuged and extracted overnight with shaking at 100 rpm. For samples in starvation, extraction was scaled-down to 50 µL of solvent and ∼50 mg of grounded tissue. Each extract was then vortexed and centrifuged, and the supernatant was transferred into glass vials for gas chromatographic analyses.

Constituents in hexane extracts were identified by GC-MS and quantified by GC-FID. The GC-MS system comprised a Gerstel 2.5.2 autosampler, a 7890A gas chromatograph and a 5975C quadrupole MS (Agilent Technologies, Santa Clara, USA). Aliquots of extracts (1 μL) were injected in splitless mode and the MS was adjusted to manufacturer’s recommendations using tris-(perfluorobutyl)-amine (FC-43). We used the following MS source conditions: injection temperature 250°C, transfer line 280°C, ion source 230°C, quadrupole 150°C, 70 eV (EI mode), 2.66 scans s^-1^ and scanning range *m*/*z* 50-600. The GC-MS column was a VF-5MS (30 m × 250 μm i.d. with 0.25 μm film thickness) fitted with a 10 m EZ-Guard column (J & W, Agilent). Helium was used as the carrier gas at a flow rate of 1 mL min^-1^. The column temperature was held at 70°C for 6 min following injection, then ramped at 10°C min^-1^ to 320°C and held at that temperature for 5 min. Mass spectra were evaluated using Agilent MSD ChemStation (E.02.02.1431) and sesquiterpenes were identified using a mass spectra library described before [94]. Other constituents were identified using the NIST 11 database (https://chemdata.nist.gov/dokuwiki/doku.php?id=chemdata:start). Constituent identity was confirmed by their relative retention times and by comparison to authentic standards where available, including the FAME mix (Sigma-Aldrich, St. Louis, USA).

Quantification of constituents in extracts was performed using a GC-FID system comprising a Perkin Elmer Autosystem XC (Perkin Elmer, Melbourne, Australia) fitted with a low polarity Zebron ZB-5 (30 m × 250 μm i.d., Phenomenex) column. Aliquots of extracts (2 μL) were injected under a split flow of 7.5 mL He min^-1^. The injector and detector temperatures were 275 and 320°C, respectively. The temperature program was the same as for the GC-MS analyses. Quantification of constituents in each class was based on calibration series of commercial standards of d-limonene (monoterpenes and C_8_ volatiles), aromadendrene (aromadendrane sesquiterpene hydrocarbons), valencene (other bicyclic sesquiterpene hydrocarbons), caryophyllene oxide (oxygenated sesquiterpenes), hexadecane (hydrocarbons), palmitic acid (fatty acids and aldehydes), phytol (diterpenes) and stigmasterol (phytosterols; Sigma-Aldrich).

### Statistical analysis

Statistical significance was calculated using ANOVA from agricolae package using Rstudio software (https://www.rstudio.com) and corrected by Tukey HSD (alpha = 0.05) for levels calculations. In other cases, built-in packages in R were used for pairwise *t*-test, Wilcoxon rank sum test and binominal test. In each figure it is explained which test was used.

## ACKNOWLEDGEMENTS

We thank Dr. Teruyuki Niimi (National Institute for Basic Biology) and Dr. Takahisa Miyatake (Okayama University) for identification of *Armadillidium vulgare*. The support of plant cultivation rooms was provided by the Model Plant Research Facility of National Institute for Basic Biology. We thank Dr. Magnus Eklund (Uppsala University) for his help with transcriptional reporters. We thank Dr. Detlef Weigel (Max Planck Institute, Germany) for kindly providing pFK247 expression vector, Dr. Kimitsune Ishizaki (Kobe University) for the cloning vectors and Dr. Keisuke Inoue for providing pMpGE_En04 and pBC-14. We thank Dr. Saiko Yoshida and Dr. Peter Huijser for assistance with confocal imaging and image analysis and Michelle van der Gragt for GUS stainings of BoGa plants.

## AUTHOR CONTRIBUTIONS

FR, EB, JLB, MT and JEM designed the research; MT and JEM conceived the project; FR, EB, SNF, JQD, RM, TD, SZ, TK and TU performed research, FR, EB, JLB, MT and JEM wrote the paper.

## SUPPLEMENTARY INFORMATION

Table S1. Differentially expressed genes upon axenic conditions in Mp*c1hdz*.

Table S2. Differentially expressed genes upon non-axenic conditions in Mp*c1hdz.*

Table S3. GO categories for down-regulated genes in Mp*c1hdz*.

Table S4. Chemical compounds identification in hexane extracts.

Table S5. Primer list.

Table S6. Summary statistics of RNA-seq experiments.

## Supplementary Methods and Figures

### Supplementary Methods

#### Transient transformation of tobacco leaves

*Nicotiana benthamiana* leaves were syringe-infiltrated with a bacterial suspension of *Agrobacterium* (LBA4404) harboring the construct _*pro*_*35S:GFP-*Mp*C1HDZ* or water (mock) following a previously established protocol [1]. The abaxial epidermis of leaves was imaged using confocal microscopy (Leica TCS SP8 Compact, Leica Microsystems, Wetzlar, Alemania). For GFP channel an excitation at 488nm, and emission at 498-510nm; for chloroplast autofluorescence channel we used the same excitation wave length and emission at 595-669nm. Images were acquired and processed using Leica Application Suite (LAS-AF, v.2.7.3) and ImageJ software (National Institute of Health, https://imagej.nih.gov/ij/) was used for generating merged pictures.

#### RT-qPCR analysis

Total RNA was purified from 3-week-old *M. polymorpha* plants using Trizol reagent (Invitrogen, Carlsbad, CA, USA) according to the manufacturer’s instructions. 1µg of RNA was reverse-transcribed using oligo(dT)18 and MMLV reverse transcriptase II (Promega, Madison, WA, USA). Quantitative real-time PCR (qPCR) was performed using a StepOnePlus Real-Time PCR System (Applied Biosystems); each reaction contained a final volume of 20 µL that included 2 µL of SyBr green (4×), 8 pmol of each primer, 2 mM MgCl_2_, 10 µL of a 1/15 dilution of the RT reaction and 0.1 µL of Taq Polymerase (Invitrogen). Thermocycler parameters were as follows: 94ºC 12 sec, 60ºC 12 sec, 72ºC 12 sec. Fluorescence was quantified over 40 cycles at 72ºC. Specific primers for each gene were designed and are listed in Table S5. The mRNA levels were quantified by normalizing their levels to the levels of the ACTIN 7 [2] using the Ct method. All of the reactions were performed with three biological replicates. Each replicate was obtained by pooling tissue from 3 to 4 individual plants.

---

**Figure S1.**
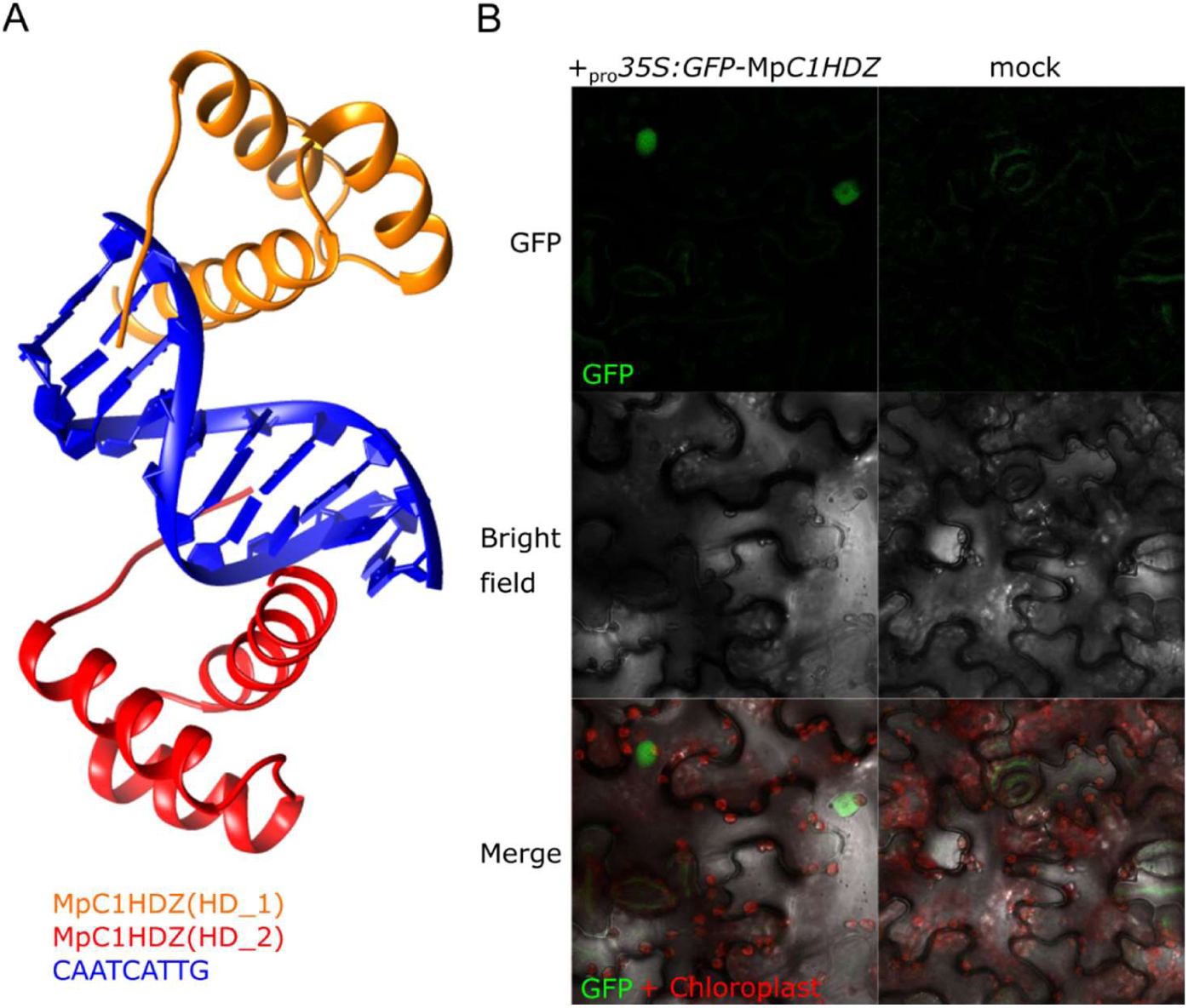
MpC1HDZ resembles structural features of angiosperms C1HDZ orthologs. (A) Ribbon representation of the 3D model of two MpC1HDZ DNA binding domains interacting with the pseudo-palindromic sequence CAATCATTG. The structure was determined using Phyre2 and based on a *Drosophila* homeobox protein (PDB: 1JGG). (B) Subcellular localization of MpC1HDZ in *Nicotiana benthamiana* leaves infiltrated with a bacterial suspension of *A. tumefaciens* harboring the construct _*pro*_*35S:GFP-*Mp*C1HDZ* or mock (water). The abaxial epidermis of rosette leaves was imaged using confocal microscopy. The photos represent the GFP specific channel, bright-field, and an overlay of both (Merge).

**Figure S2.**
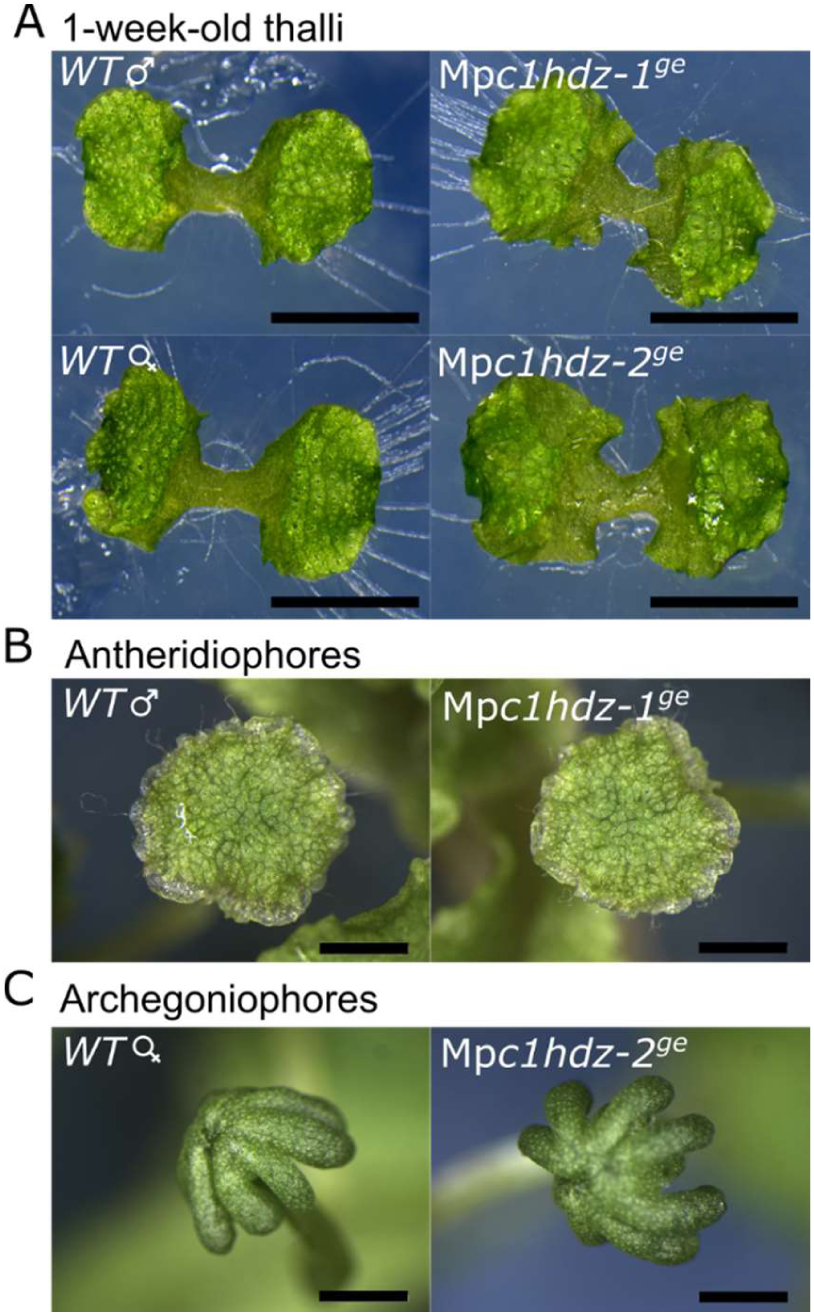
Plant morphology of wild-type and mutant alleles. Thallus of wild-type and Mp*c1hdz* plants 7 days after germination (DAG). Wild-type and Mp*c1hdz* gametangiophores; male, antheridiophore; female, archegoniophore. Scale bars (2mm).

**Figure S3.**
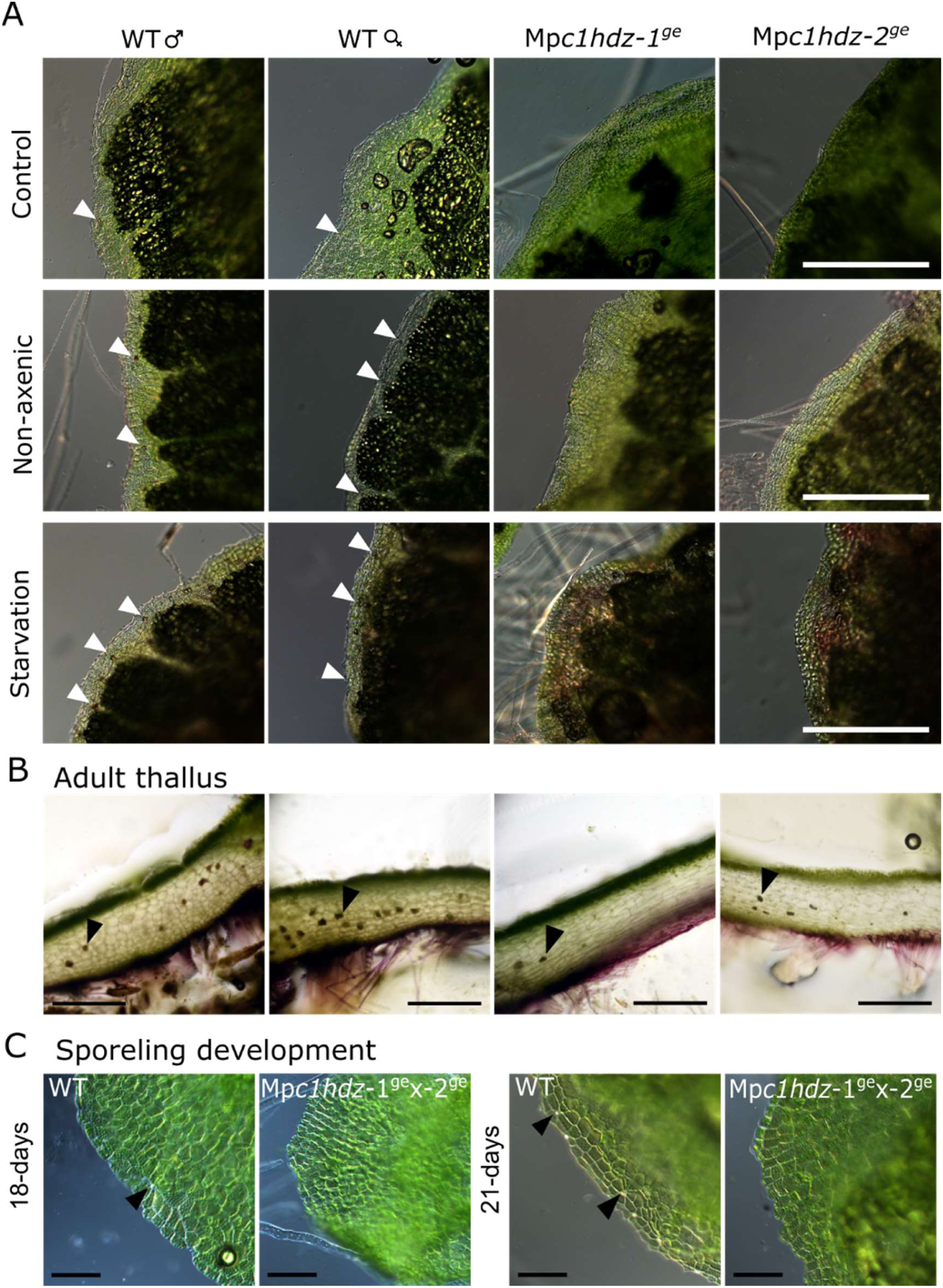
Plant growth responses upon abiotic stress. (A) Representative compound microscopy pictures of 3-week-old plants thallus grown in control conditions for one week and then transferred again to control conditions, or starvation conditions (1/100 Gamborg’s B5 medium) or non-axenic (rockwool as substrate). Arrows point to oil bodies. Scale bars (0.5mm). (B) Transverse sections of 30-day-old thallus grown in non-axenic conditions. (C) Representative pictures of wild-type and Mp*c1hdz* sporelings. Black arrow indicates oil bodies. Scale bars (0.1mm).

**Figure S4.**
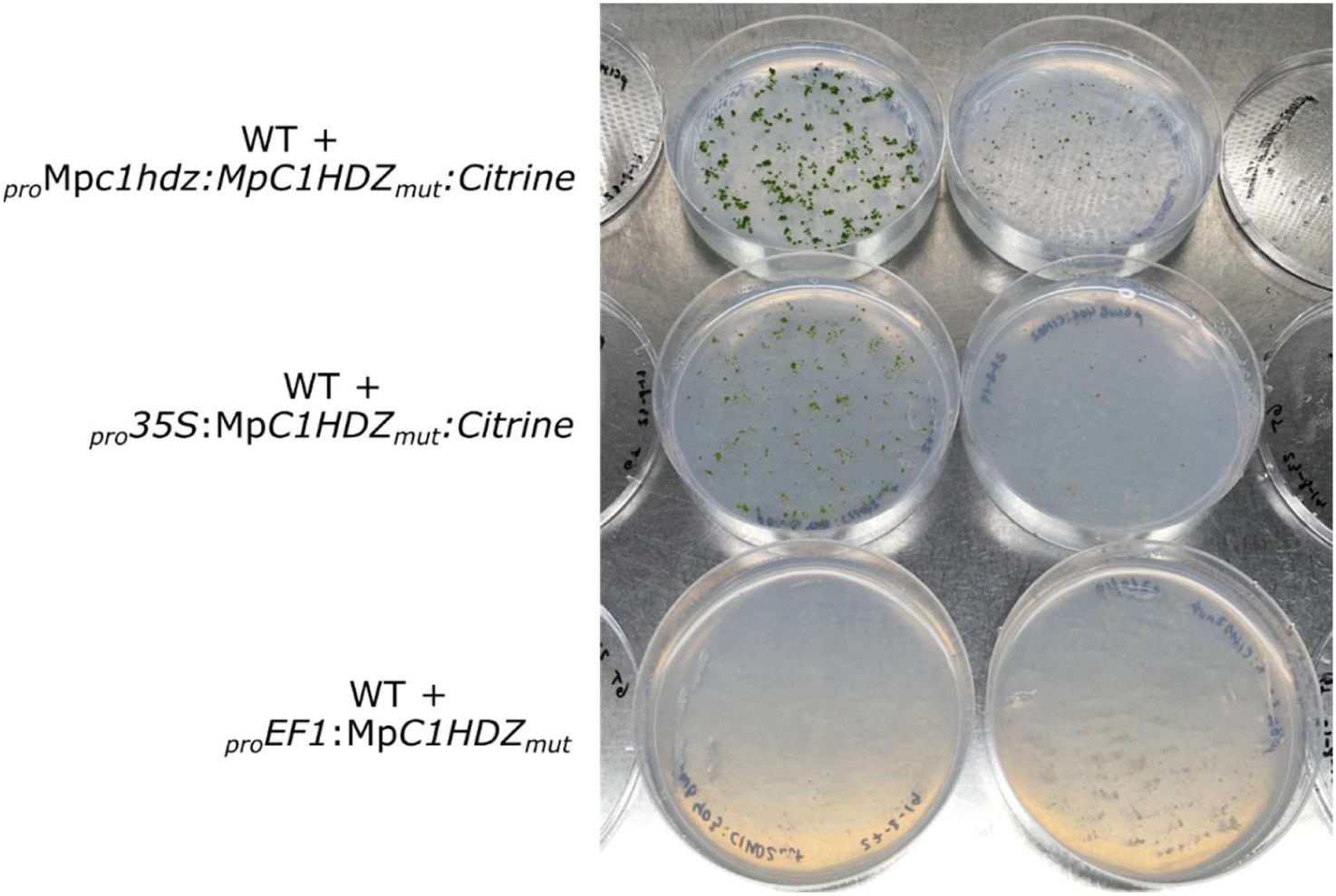
Constitutive overexpression of Mp*C1HDZ* may be lethal. Representative pictures of wild type (*WT)* spores transformed with a vector constitutively overexpressing Mp*C1HDZ*. After 10 day of selection on antibiotic plates, most plants did not survive compared to a control transformation. Two biological replicates are shown.

**Figure S5.**
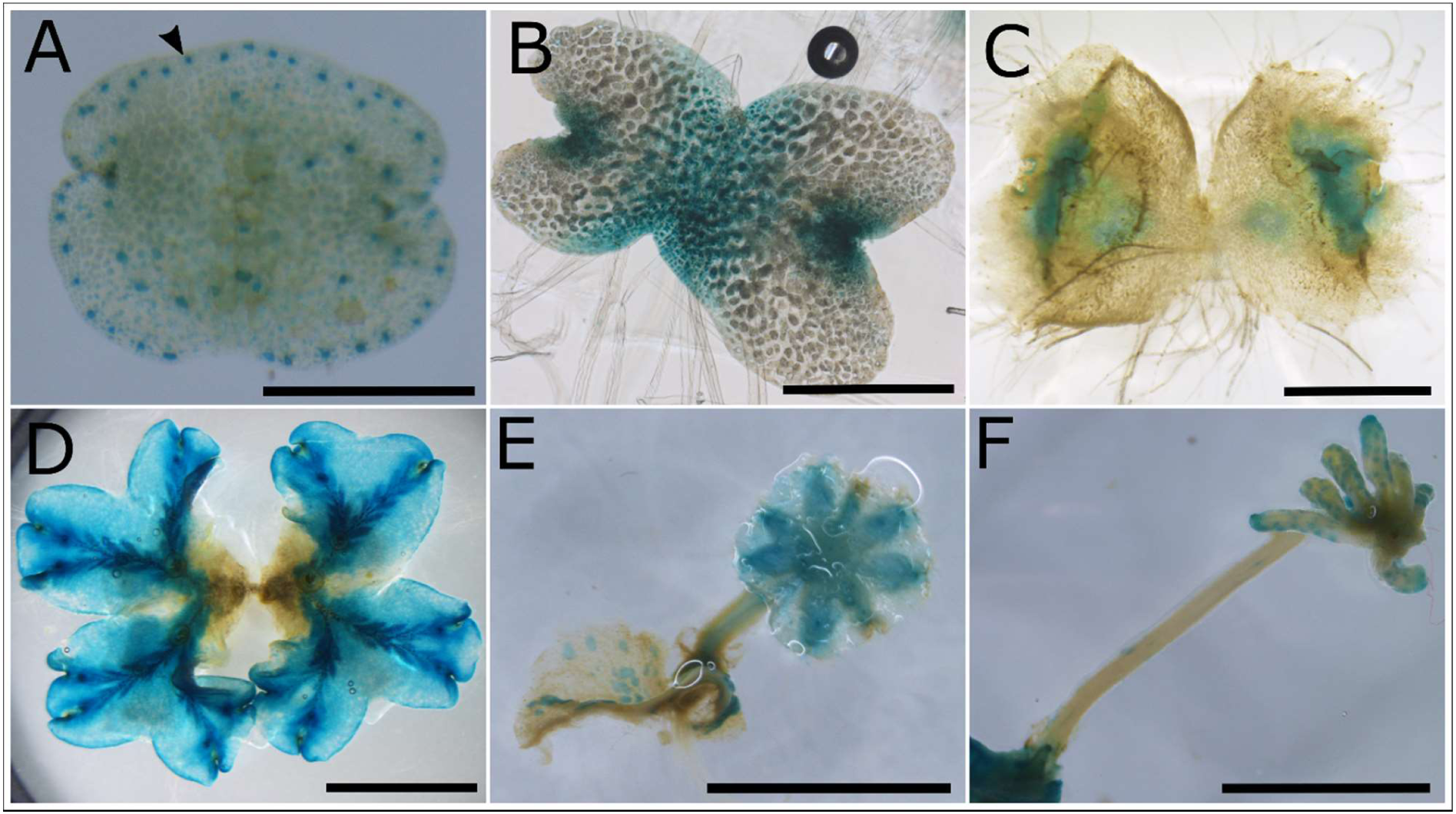
Mp*C1HDZ* promoter activity in BoGa ecotype. Mp*C1HDZ* expression pattern through different life stages visualized by β-glucuronidase staining in _*pro*_Mp*C1HDZ* ^*3kb*^:*GUS* lines (A – F). Vegetative propagule, gemma (A), 3 day-old gemmaling (B), one week old gemmaling (C), three weeks old mature thallus (D), male sexual structure, antheridiophore (E) and female sexual structure, archegoniophore (F).

**Figure S6.**
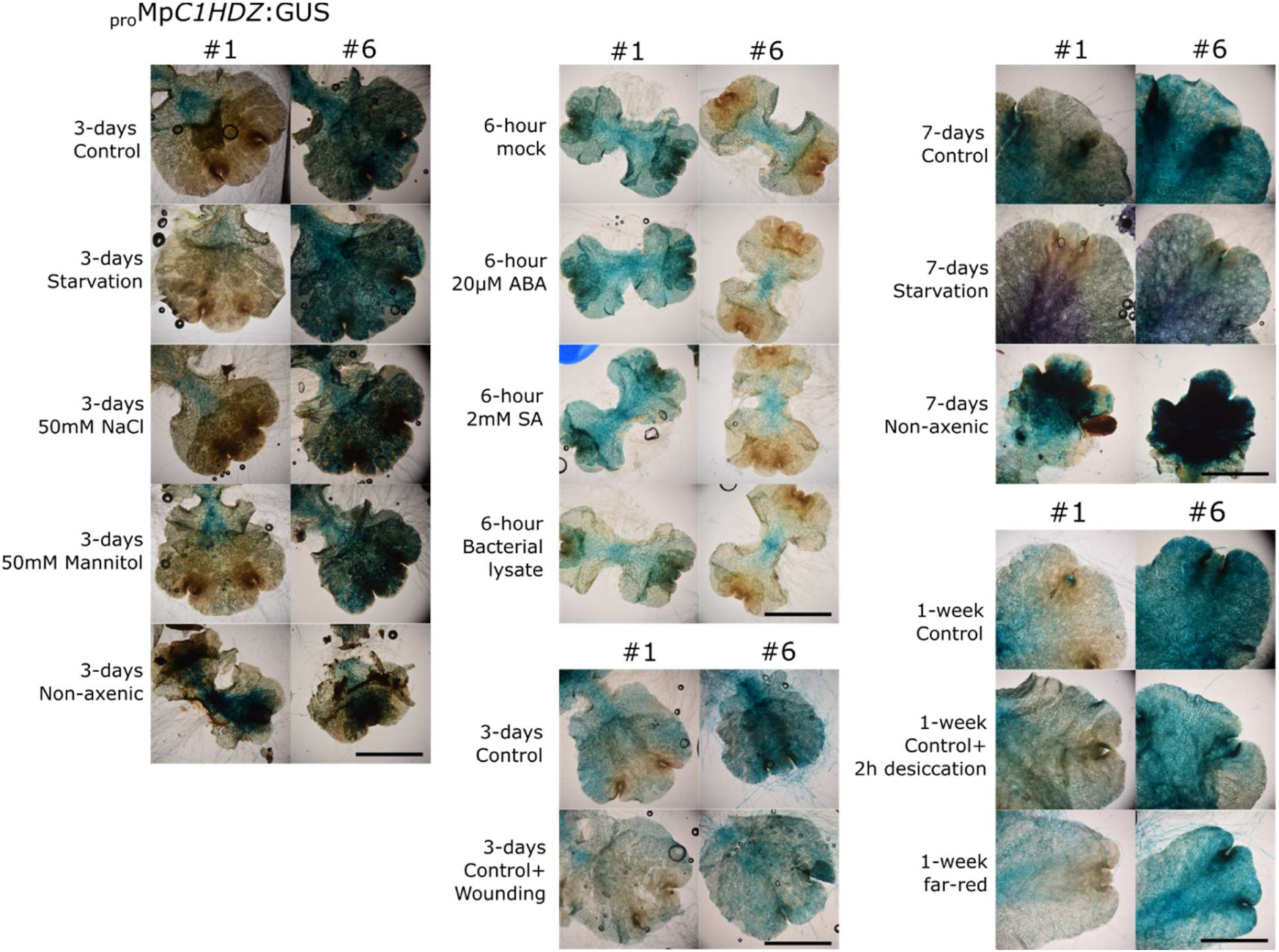
Mp*C1HDZ* promoter activity upon abiotic stress treatments. GUS staining of _*pro*_Mp*C1HDZ*^*4.6kb*^:*GUS* transcriptional reporter under different treatments. Plants were grown for one week and then transferred to new plates containing different media and treatments as indicated in the left. Scale bars represent 0.5mm.

**Figure S7.**
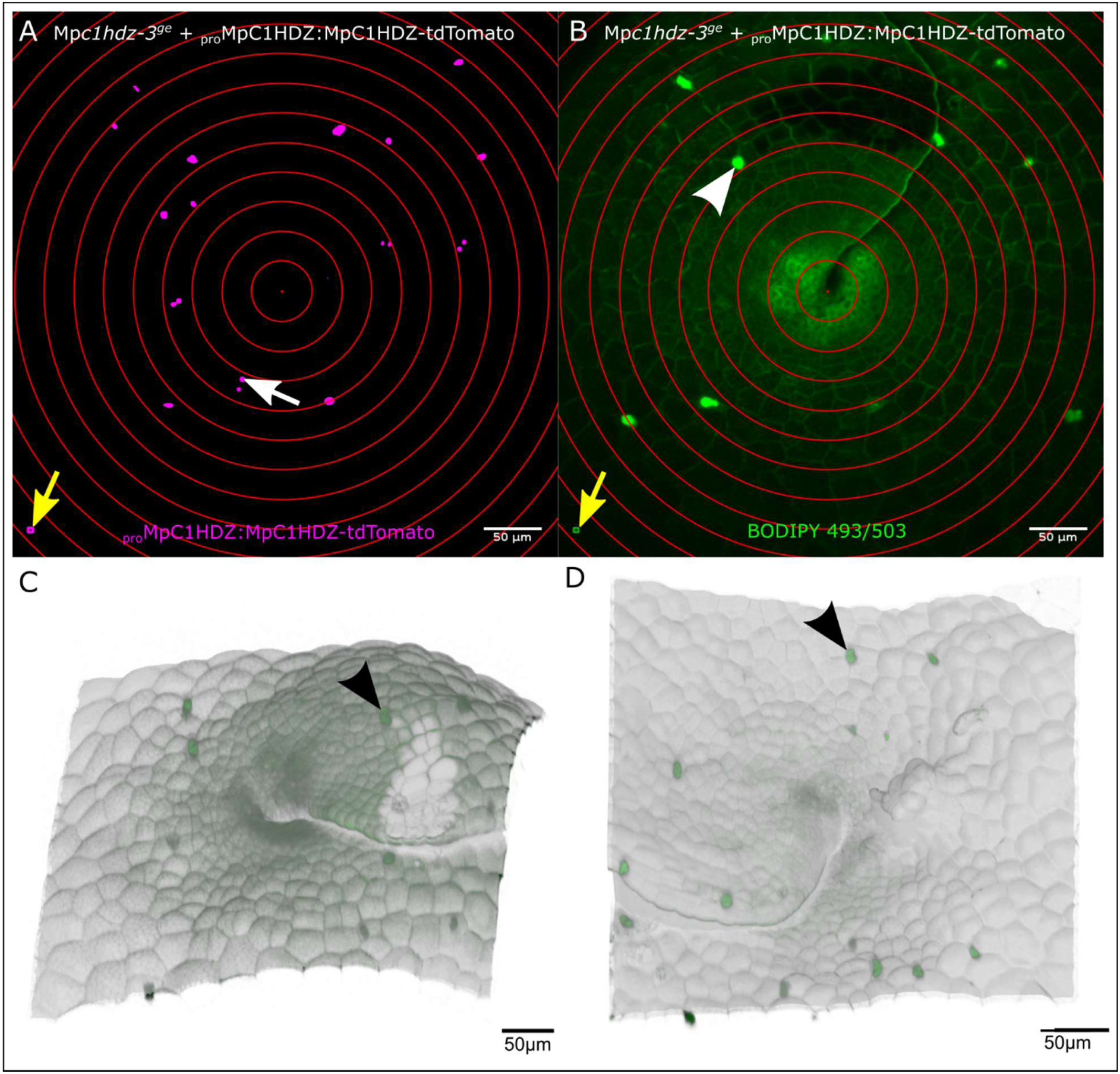
Measurement of distance from apical notch to nuclei highly expressing MpC1HDZ and oil bodies. Apical notch area of Mp*c1hdz-3* gemmaling complemented with _*pro*_Mp*C1HDZ*^*3kb*^:Mp*C1HDZ-tdTomato* and stained with BODIPY 493/503 dye. (A, B) 2D projection of a representative confocal z-stack from Fig. 2G with concentric circles used for quantification in Fig. 2H. MpC1HDZ-tdTomato channel with nuclei above the threshold (A) and BODIPY 493/503 channel with oil bodies (B). Concentric circles are centered in the middle of the apical notch and are interspaced by 25 μm. White arrow points to a bright nucleus, arrowhead points to an oil body, and yellow arrows point to the pixels used for calibration of “sum slices” z-stack projection. (C, D) 3D representation of apical notch surface from two replicates with BODIPY signal 493/503 shown in green. Arrowheads point to oil bodies. The sample in C corresponds to A and B. Note that the circles in A and B represent approximate measure because of variability in apical notch area 3D morphology (C, D). Scale bars represent 50 μm.

**Figure S8.**
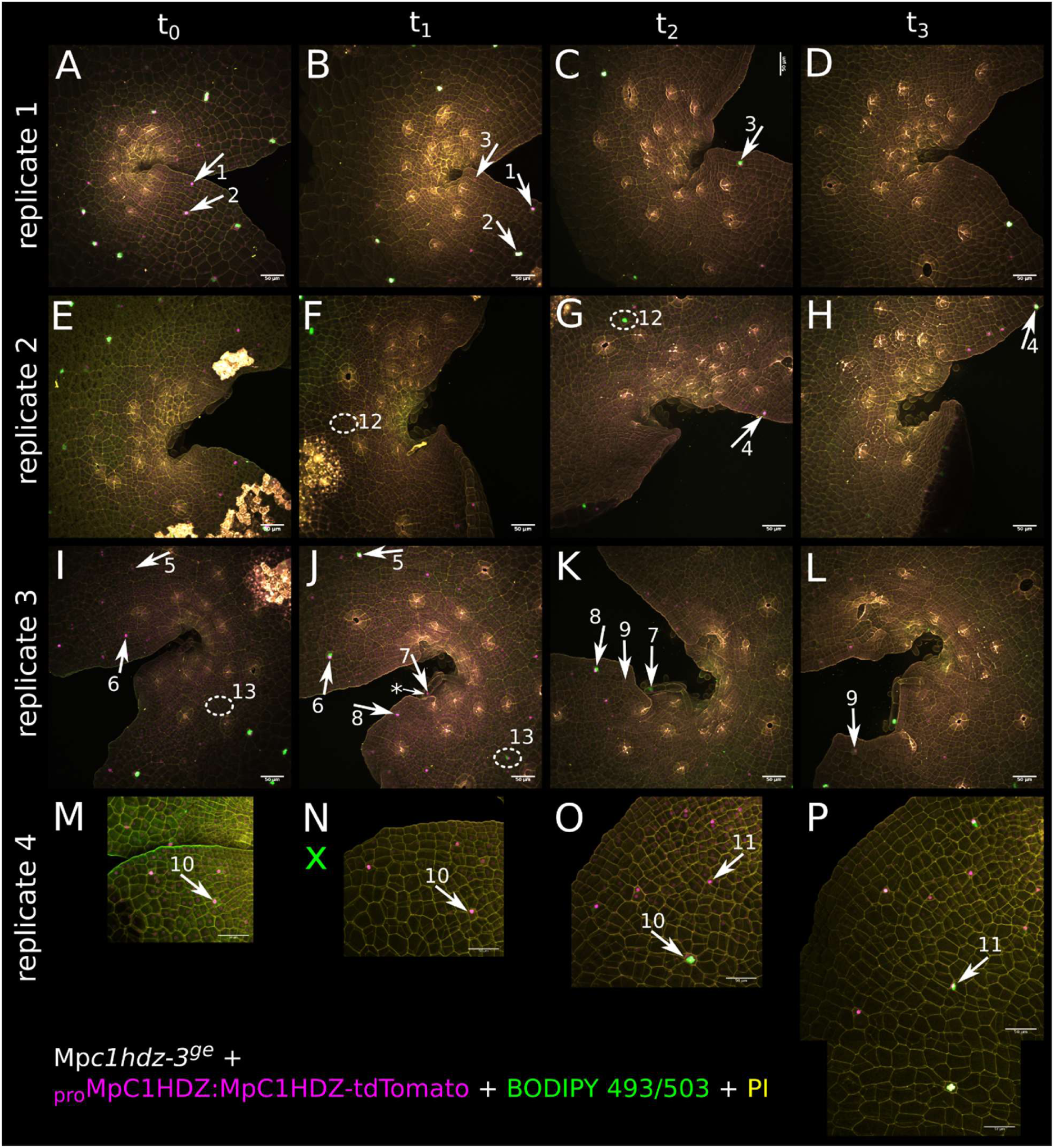
Overview of oil body cell tracking time lapse. Time series of oil body cell differentiation in apical notch area of 4 day-old Mpc1hdz-3 gemmalings complemented with proMpC1HDZ3kb:MpC1HDZ-tdTomato (magenta) and stained with BODIPY 493/503 (green) and PI (yellow) directly before CLSM imaging. The same area was followed for 48 h in 12 h intervals (t0 – t1, shown from left to right). Standard deviation projections of confocal z-stacks of four replicates are shown: replicate 1 (A – D), replicate 2 (E – H), replicate 3 (I – L) and replicate 4 (M – P). All newly-formed oil body cells are equivalently numbered and marked before and after oil body formation event. Newly-formed oil body cells that had exceptionally high MpC1HDZ-tdTomato expression compared to surrounding cells in previous time point are indicated with arrows (numbers 1 – 11). Newly-formed oil body cells that did not have outstandingly bright MpC1HDZ-tdTomato signal in previous time point are marked with dashed circles (numbers 12 and 13). The latter observations may suggest that the time interval used for observation is longer than the minimal period of high MpC1HDZ expression needed for oil body formation. Alternatively, high MpC1HDZ expression may not be necessary for oil body formation in a small proportion of cells, which would also explain presence of a few oil bodies in Mpc1hdz mutants (Figure 1, Figure S3). Asterisk and a small arrow mark a shadow observed in cell 7 at the spot stained with BODIPY in the next time point. This may indicate that this cell was developing an oil body, but did not yet accumulate the compounds stainable with BODIPY by the time of observation. This was also the only observed oil body formation in a large cell that most likely belongs to a ventral scale. One case where BODIPY was not applied is marked with green X. Signal intensities between the images are not comparable due to varying z-stack sizes. Scale bars represent 50 μm.

**Figure S9.**
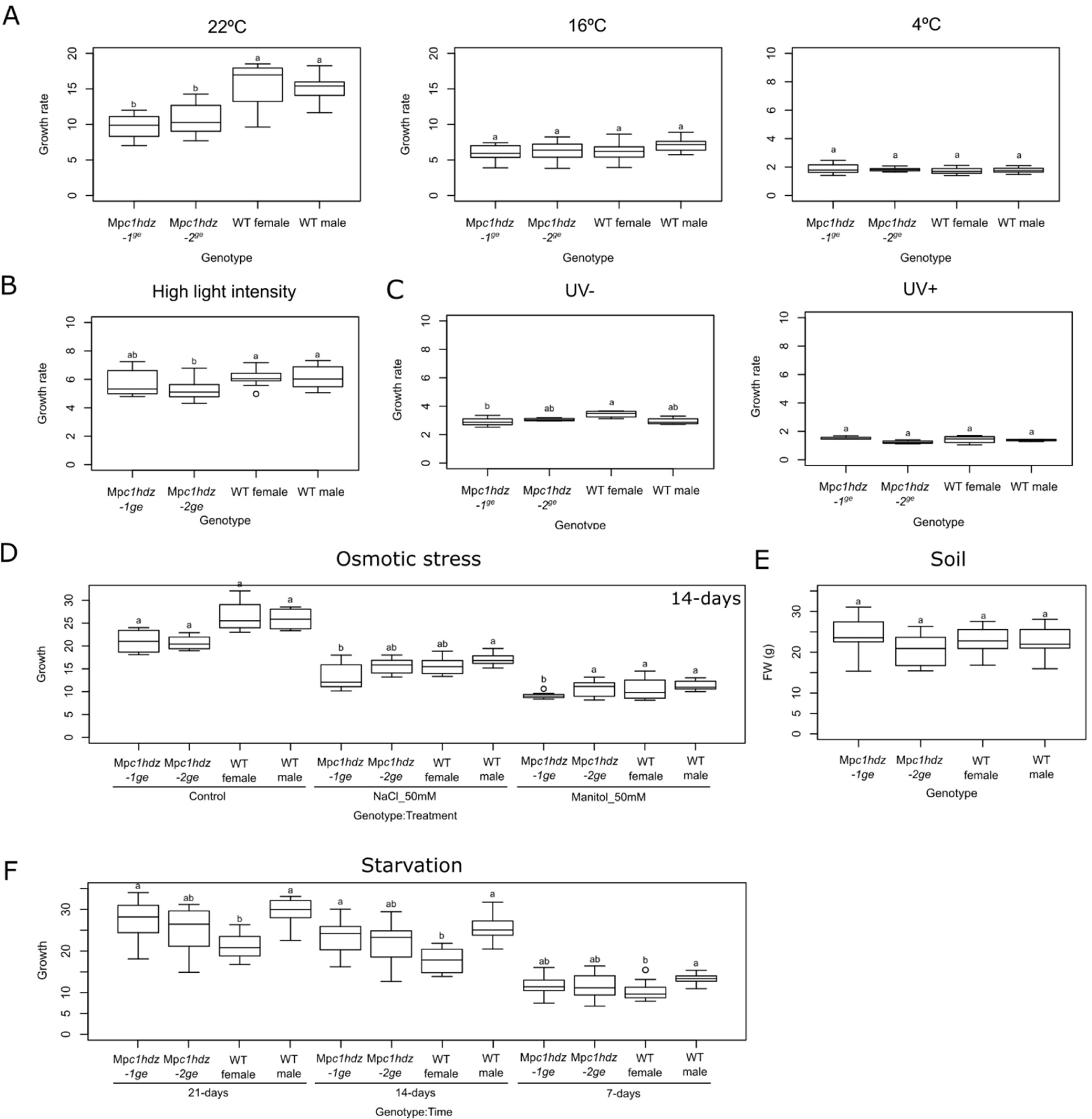
Thalli growth rate of Mp*c1hdz* under different abiotic stress treatments. (A-F) Phenotypic effect of abiotic stress treatments on wild-type and Mp*c1hdz* plants. Plants were grown axenically on agar plates with 1/2 strength Gamborg B5 media for 1-week and then transferred to a new plate. Growth was measured as lamina exposed area compared to the area after 1 week of treatment or different time-points. Cold treatment was performed at 4ºC; high light intensity (HLI) was performed using 1000 µmol m^-2^s^-1^LED lights; UV treatment was performed in a laminar flow bench UV-C light for 1 hour and recovered for an additional week under control conditions. For osmotic stress, media was supplemented with 50 mM NaCl or mannitol, grown for 2-weeks (in Figure 1 it is shown after 1-week). For starvation treatments, plants were placed in 1/100 Gamborg B5 media. For non-axenic growth, plants were grown on pots in a growth chamber using rockwool as a substrate. Statistical difference was tested using ANOVA followed by Tukey HSD *p.*value < 0.05; letters indicate statistically significant groups. Representative pictures of each treatment are shown.

**Figure S10.**
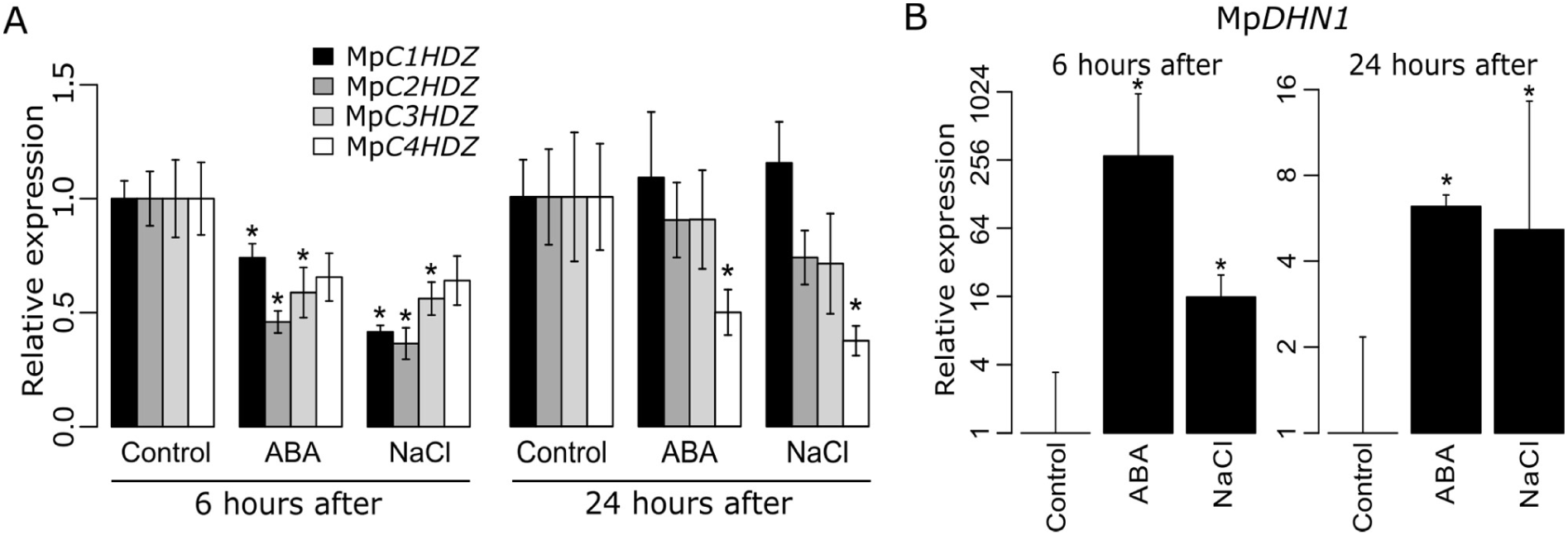
Expression of *HDZ* genes under ABA and abiotic stress treatments. (A) RT-qPCR analysis of all *HDZ* genes in *M. polymorpha* under ABA (20 uM) and NaCl (200 mM) treatments. RNA samples were taken after 6 and 24 h. (B) The abiotic stress marker gene (Mp*DHN1*) was used as a treatment control. Statistical differences are marked as asterisks and computed pairwise *t*-test (*p*.value < 0.05) comparing each treatment to the corresponding control.

**Figure S11.**
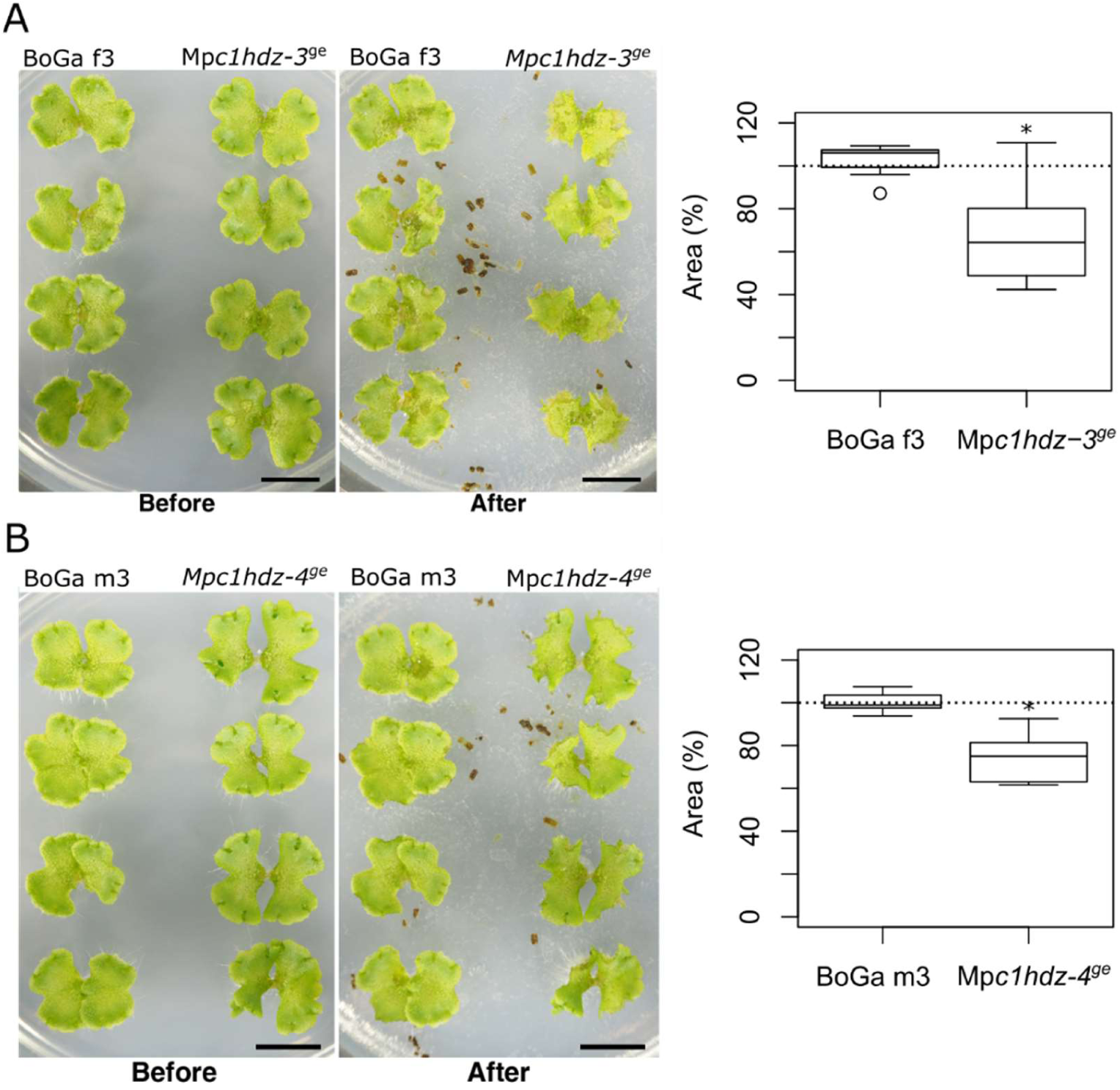
Pill bug feeding assay on mutant alleles of the BoGa ecotype. Starved pill bugs and 13-day-old thalli (Before) were co-cultivated for 24 hours (After). Female wild-type (BOGA f3) and Mp*c1hdz-3*^*ge*^ mutant thalli or male wild-type (BOGA m3) and Mp*c1hdz-4*^*ge*^ mutant thalli were used in the experiments. Bars = 1 cm. Thallus area change indicates the difference between thallus area (mm^2^) of “After” and that of “Before.” Thallus change ratio indicates the ratio of thallus area of “After” to that of “Before” (n = 12 thalli for each genotype). Error bars indicate the mean ± SD. Statistical analysis between wild-type and mutant plants was conducted by a two-tailed Welch’s *t*-test.

**Figure S12.**
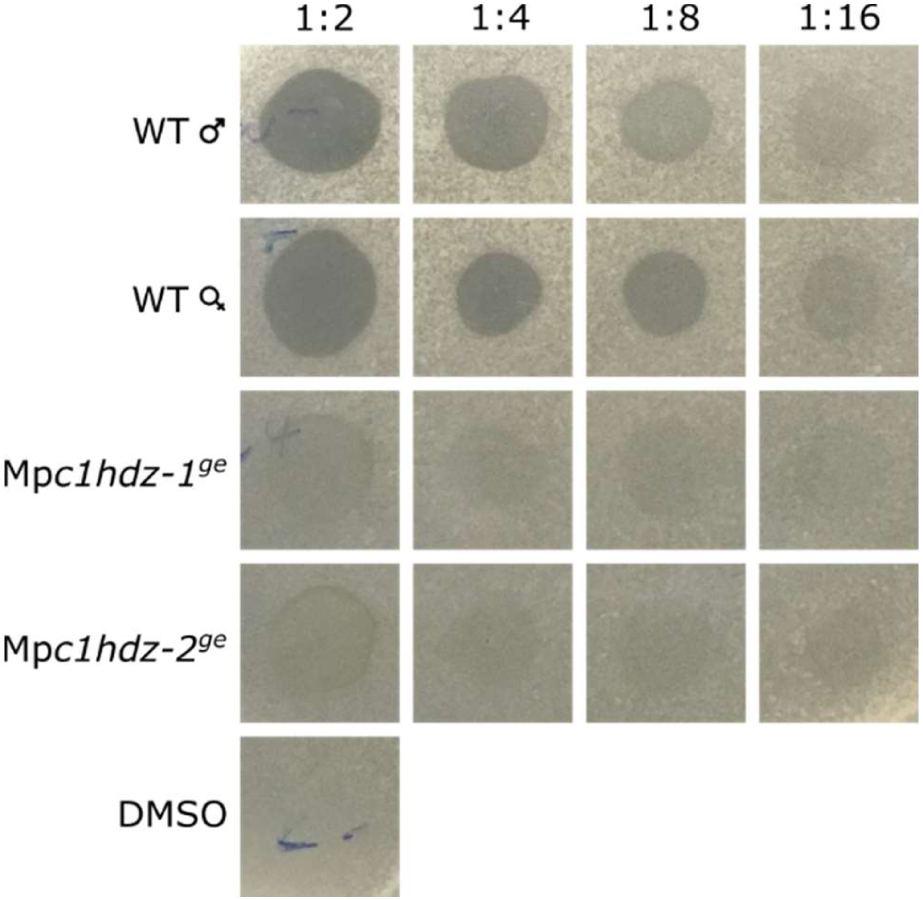
Dose-dependence effect of plant extract on the growth of *Bacillus subtilis*. Serial dilution of *Bacillus subtilis* overlay assay using ethanol extracts in DMSO from wild-type and Mp*c1hdz* mutant thalli.

**Figure S13.**
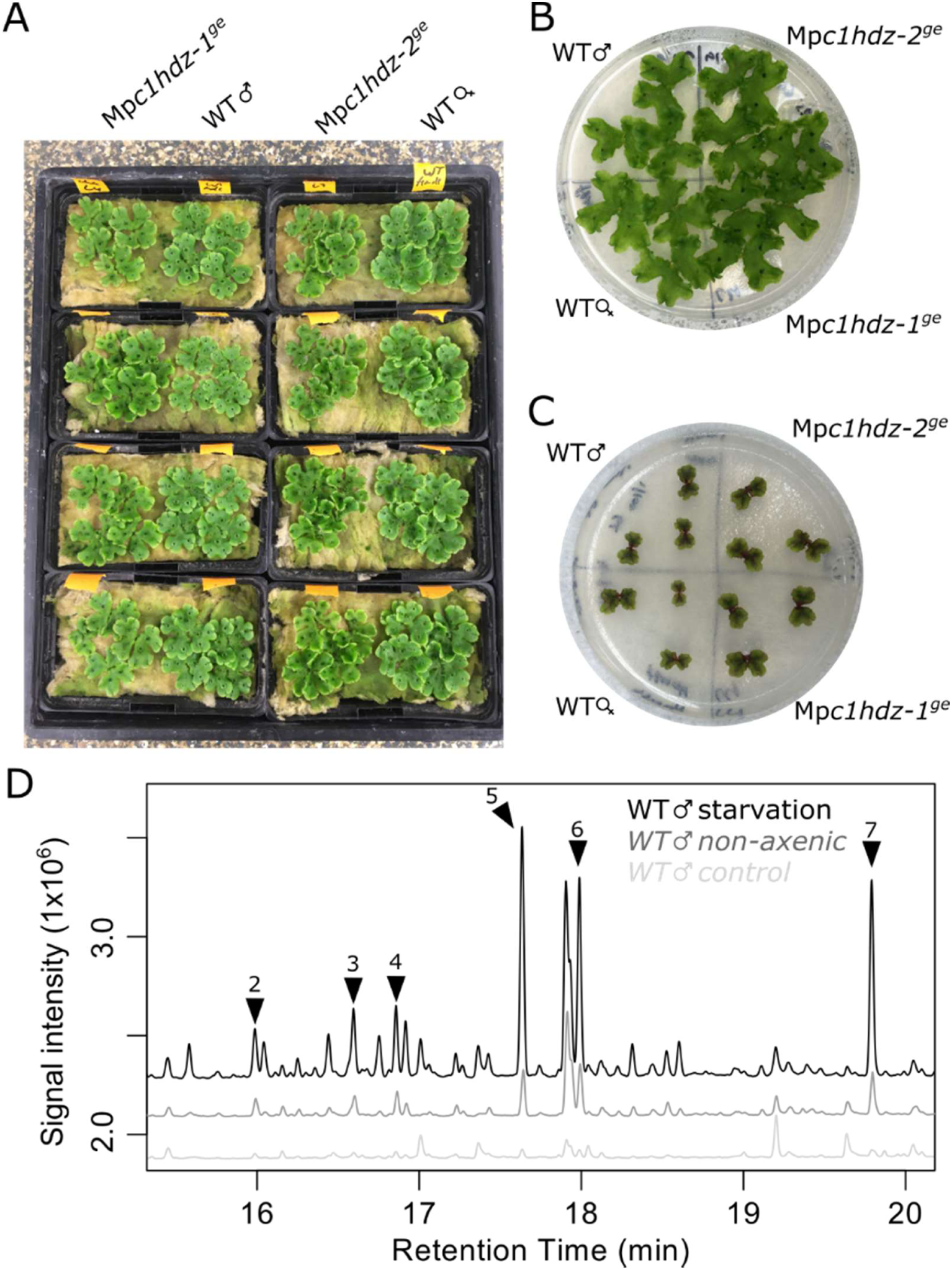
Representative figures of plants after treatments. (A) 3-week old wild-type and Mp*c1hdz* plants growing in non-axenic conditions (rockwool) (B) 3-week old wild-type and Mp*c1hdz* plants growing in axenic conditions 1/2 Gamborg B5 (control). (C) 3-week old wild-type and Mp*c1hdz* plants growing in starvation treatments 1/100 Gamborg B5.

**Figure S14.**
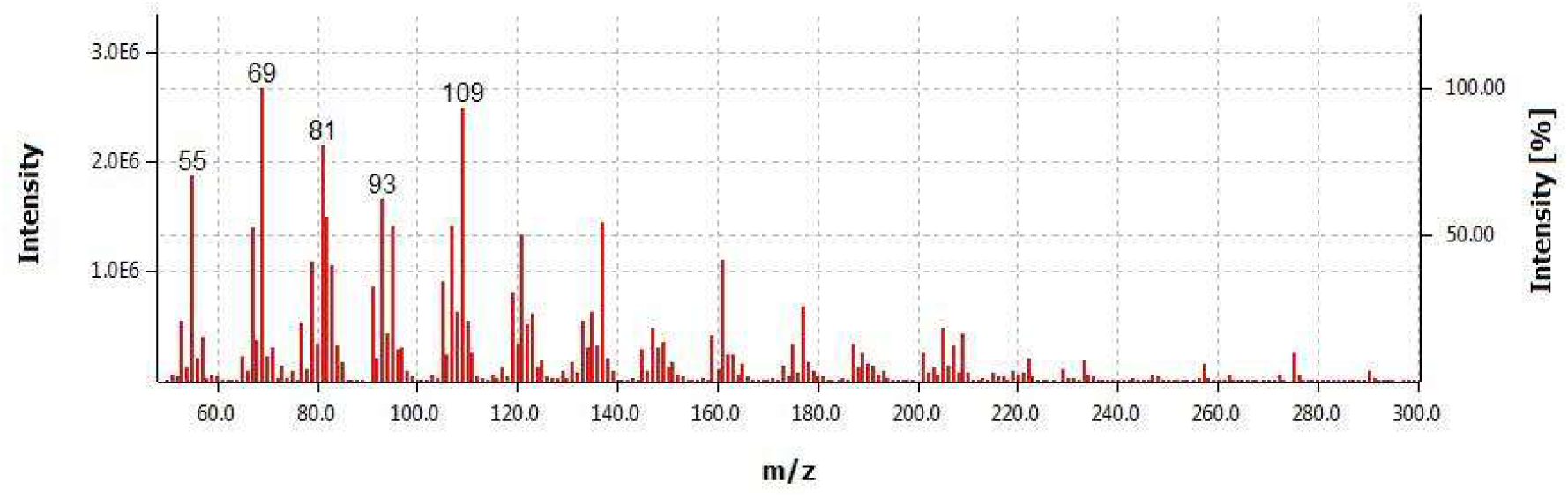
Mass spectrum of unidentified compound U13. Mass spectrum of peak 10 corresponding to the unidentified compound U13. Based on MS profiles of NIST library, we tentatively propose it is a terpenoid.

